# Monocyte Migration Emerges from a Divergent Chemokine Signaling Network

**DOI:** 10.64898/2026.04.29.721539

**Authors:** Sean S. So, Alexis Lona, Rina Pokhrel, Alexandra L. Morgan, Mila Saltikova, Thy Nguyen, Willow Carretero Chavez, Tony Ngo, Harley R. Robinson, Cheng Huang, Shankar Raj Devkota, Ram Prasad Bhusal, David H. Drewry, Joel R. Steele, Ralf B. Schittenhelm, Tracy M. Handel, Simon R. Foster, Irina Kufareva, Martin J. Stone

**Author notes:** Correspondence: SRF; IK; MJS. These authors contributed equally: Sean S. So, Alexis Lona, Rina Pokhrel.

## Abstract

Migration of leukocytes in the context of immune homeostasis or inflammatory diseases is regulated by activation of chemokine receptors by chemokine ligands. To elucidate how these interactions give rise to cell migration, we mapped the chemokine-stimulated signal transduction network in monocytic THP-1 cells. Global phosphoproteomics revealed 630 time-resolved changes in phosphorylated proteins downstream of the chemokine receptor CCR2. We used the “PHONEMeS” network modeling algorithm to generate the most parsimonious signal transduction network consistent with the observed protein phosphorylation data. The CCR2 signaling network is highly divergent, acting via multiple branches to regulate proteins required for cell migration. We validated this model using kinase inhibitors targeting different branches of the network and successfully blocked chemokine-stimulated cell migration. Thus, chemotaxis is an emergent property resulting from an integrated cellular response to divergent signaling pathways. This paradigm suggests that physiological regulation or pharmacological blockade of chemokine-driven inflammation could potentially be achieved by inhibiting any of the divergent pathways within the network.

## Introduction

Inflammation may be an acute homeostatic response to injury or infection leading to tissue protection and repair, or a pathological state associated with chronic tissue damage and dysfunction. In both cases, inflammation is characterized at the cellular level by the recruitment of circulating leukocytes into the affected tissues.

Leukocyte migration is orchestrated by the secretion of chemokines within tissues and their interactions with chemokine receptors expressed on leukocytes^1–4^. Selective inhibition or genetic deletion of chemokine receptors suppresses inflammation *in vivo*^5–8^, but translation of these outcomes to the clinic has been challenging^7–9^. In particular, antagonists of CC chemokine receptor 2 (CCR2) inhibited monocyte recruitment in preclinical models of atherosclerosis and rheumatoid arthritis but were not efficacious in human clinical trials for these diseases^8,10–13^.

Chemokine receptor activation leads to changes in actin cytoskeleton dynamics and coordinated activation/inactivation of adhesion molecules, resulting in directional cell migration^14^. However, the network of signaling events leading from chemokine receptor activation to these cellular changes remains only partly defined^15–18^. Identifying critical pathways within the network will present tractable opportunities for pharmacological intervention in inflammatory diseases.

Cellular signaling networks involve coordinated cascades of protein phosphorylation. Major advances in mass spectrometry-based phosphoproteomics have enabled global identification of thousands of proteins involved in specific signaling responses^19^. Importantly, phosphoproteomics data constitute both direct evidence for kinase and phosphatase activity as well as indirect evidence for unobserved signaling events on which these phosphorylation events are dependent. Thus, a powerful and unbiased approach for assembling signaling networks is to interpret phosphoproteomics data in the context of prior knowledge of relationships between signaling molecules^20–23^.

Here, we define the signaling network resulting from activation of endogenous CCR2 in a monocyte cell line. Specifically, we used time-resolved phosphoproteomics data following CCR2 activation to generate a global signaling network consistent with the experimentally-observed phosphorylation events. Network analysis revealed divergent pathways regulating actin remodeling, cell polarization and adhesion. Pharmacological inhibition of these pathways suppressed chemokine-stimulated cell migration. Our results establish a paradigm in which chemokine-driven monocyte migration emerges from the temporal coordination and integration of divergent signaling pathways, and demonstrates the potential of targeting these pathways to suppress leukocyte migration as a strategy for treating inflammatory diseases.

## Results

### Global Phosphoproteomics Reveals Extensive CCR2-mediated Phosphoprotein Signaling in Monocytes

THP-1 cells are a human monocytic cell line whose chemokine receptor expression profile mirrors that of classical human monocytes (**Supplementary Fig. 1**)^24^. The chemokines CCL2 and CCL5 stimulate THP-1 cell migration and ERK1/2 phosphorylation exclusively via the chemokine receptors CCR2 and CCR1, as these responses are completely inhibited by CCR2- and CCR1-selective antagonists INCB3344 and BX471, respectively (**Fig. 1a,b**). Thus, THP-1 cells are an attractive model cell line to interrogate chemokine receptor-specific signaling relevant to human classical monocytes.

**Figure 1:**
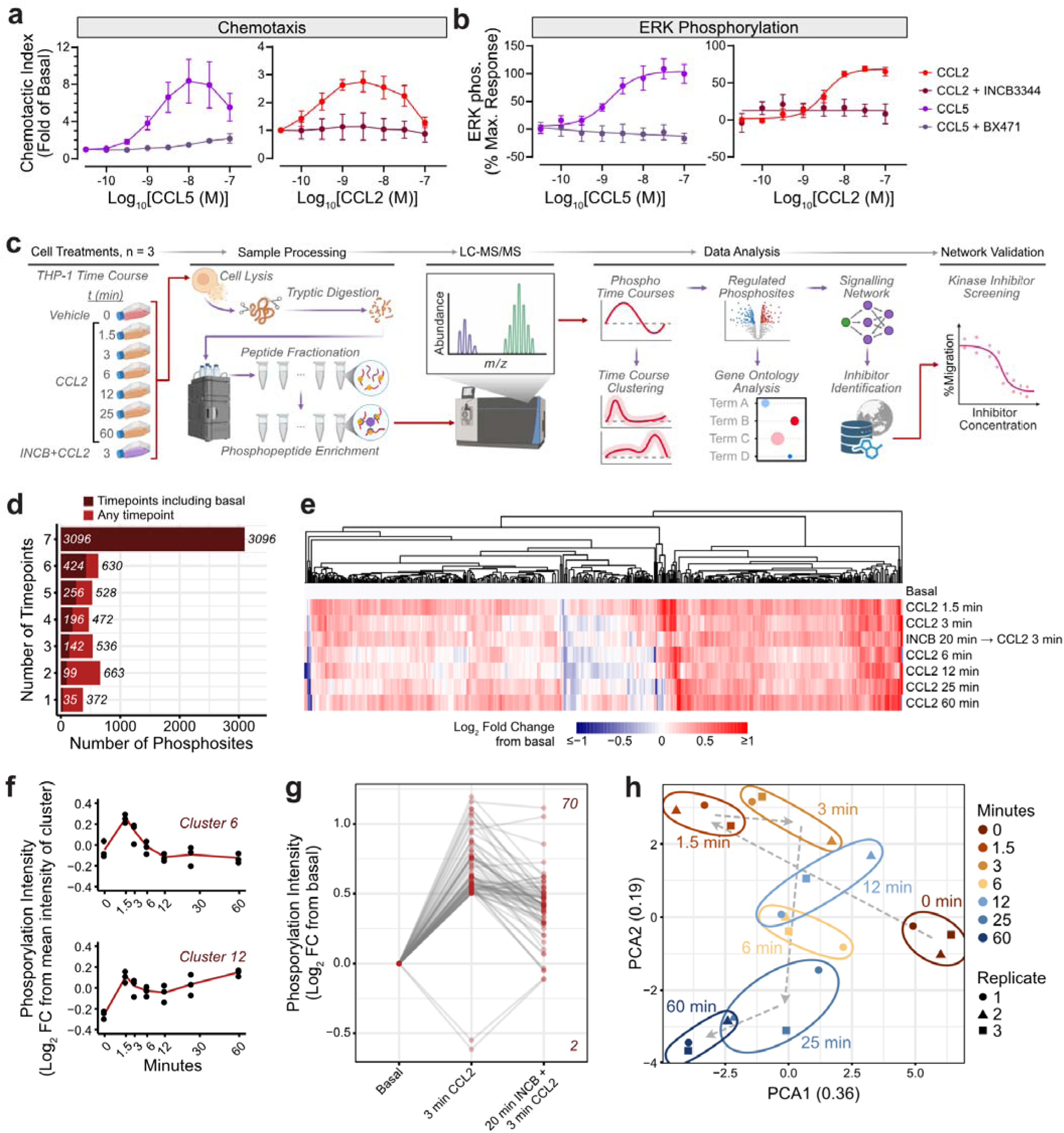
Activation of CCR2 causes extensive changes in protein phosphorylation in THP-1 monocytes. **a,b**, THP-1 cell (**a**) chemotaxis and (**b**) ERK phosphorylation in response to increasing concentrations of CCL2 (red) or CCL5 (purple). CCL2 and CCL5 signals are inhibited by 20 min pretreatment with 1 μM of INCB3344 and BX471, antagonists for CCR2 and CCR1, respectively. Points and error bars represent mean and SEM of n=4-6 experiments, performed in triplicate. **c**, Flowchart depicting the experimental and analytical pipeline in this study. **d**, The number of high quality phosphosites detected in THP-1 cells across various timepoints post CCL2 stimulation. Dark red, phosphosites detected at timepoints including basal; light red, phosphosites detected at any timepoints. **e**, Heatmap indicating the log_2_ fold-change (FC) from basal for 630 significantly varied phosphosites across various timepoints. Tile colors (blue/white/red) indicate the log_2_FC relative to basal. For clarity, the range -1.4 to -1 and 1 to 1.8 were trimmed to -1 and 1, respectively. **f**, Time-dependent log-intensity profiles of representative phosphosite clusters (Clusters 6 and 12, see **Supplementary** Fig. 4 for a full list of cluster intensity profiles). Points represent averages of 3 replicates across all cluster members. The red line indicates the mean of the replicates. Time is represented on a square-root axis. **g**, Pretreatment of THP-1 cells for 20 min with 1 μM INCB3344 prior to a 3-minute stimulation with 10 nM CCL2 inhibits most of the strong phosphorylation and dephosphorylation events. Points connected by lines indicate matched phosphosites; y-axis represents mean log_2_ FC from basal phosphorylation intensity; points are shown for all 72 phosphosites (70 upregulated, top-right; 2 downregulated, bottom-right) with |log_2_FC|>0.5 that were detected in both the absence and presence of INCB. **h**, Principal component analysis (PCA) of triplicate measurements (circles, triangles, squares) for all 630 significantly regulated phosphosites, as determined by ANOVA.

We used global phosphoproteomics^25,26^ to identify the changes in protein phosphorylation following CCR2 activation (**Fig. 1c**). THP-1 cells were stimulated with a saturating concentration of CCL2 (100 nM) for varying durations over a 1 hour period and compared to untreated cells. A 3-min CCL2 treatment of INCB3344-pretreated cells was also included to assess the CCR2-specificity of the response. Phosphorylation of cellular proteins was determined by label-free, quantitative mass spectrometry (MS) of derived phosphopeptides.

Analysis of the MS data identified 12,282 phosphosites from 2,577 proteins below a false discovery rate (FDR) cut-off of 1% in at least one sample. To improve the robustness of our interpretation of the time-resolved THP-1 phosphoproteome, we restricted our analysis to only those phosphosites that were represented in at least 2 of 3 replicates in at least one condition (excluding INCB-pretreated samples) and had localization probabilities of at least 75%. This refined data set included 6,297 phosphosites located on 2,115 proteins (**Supplementary Fig. 2a**). Among these, 3,096 phosphosites were observed at all seven time points, indicative of a high level of data completeness (**Fig. 1d**).

To identify time-resolved perturbations in phosphorylation signatures, we performed a one-way ANOVA on phosphosite intensities. We observed significant changes in phosphorylation (adjusted p-value ≤ 0.05) for 630 phosphosites, belonging to 400 proteins, within 60 min of CCL2-stimulated activation of CCR2 (**Fig. 1e; Supplementary Fig. 2b; Supplementary File 1**). Kinase Enrichment Analysis (KEA3)^27^ of these proteins indicated the enrichment of substrates of casein kinases CSK21 and CSK22, MAP kinases MK03 and MK01 (ERK1/2), cyclin-dependent kinase 9 (CDK9), and cell division cycle- (CDC-) like kinases CLK1 and CLK2, among others (**Supplementary Fig. 3**). In pairwise comparisons, 1,387 phosphosites (in 745 proteins) were significantly altered relative to the basal state (adjusted p-value < 0.05; moderated t-test with Benjamini-Hochberg correction).

Hierarchical clustering of temporal site phosphorylation profiles revealed 20 dominant patterns of CCL2-induced changes (**Supplementary Fig. 4**). In most cases, the phosphorylation spiked rapidly (significantly increasing at 1.5 min and/or 3 min, relative to the basal state) and then either quickly returned to the basal level (fewer sites; e.g., cluster 12) or persisted for up to 60 min (the majority of sites; e.g., cluster 6) (**Fig. 1f**). Cell pretreatment with INCB3344 did not completely block phosphorylation changes measured at 3 min post CCL2 stimulation (**Supplementary Fig. 2b**), presumably due to incomplete receptor occupancy; however, the strongest CCL2-induced changes (|log2FC|≥0.5) were almost universally suppressed by INCB3344, supporting CCR2 specificity (**Fig. 1g**). Principal component analysis (PCA) of significantly regulated sites reiterated these predominant variation patterns following CCL2 stimulation (**Fig. 1h**).

### Many CCL2-Regulated Phosphoproteins are Involved in Cell Migration

To identify the dominant biological functions associated with regulated phosphoproteins, we performed Gene Ontology (GO) overrepresentation analysis^28–30^ (**Fig. 2a; Supplementary File 2**). This highlighted the prevalence of proteins already known to be involved in actin cytoskeletal remodeling and/or leukocyte (although not necessarily monocyte) migration, including: actin itself, unconventional myosins, intermediate filament proteins, and direct or upstream regulators of actin filament organization, microtubule organization, cytoskeletal organization, cell polarity, stress fiber, filopodium and/or lamellipodium assembly, and adhesion (**Fig. 2b**; **Supplementary File 3**). These proteins included Rho guanine nucleotide exchange factors (ARHGEFs), Rho GTPase activating proteins (ARHGAPs), SRC kinase, and p21-activated kinases (PAKs). The majority of these proteins displayed rapid changes in phosphorylation, consistent with the temporal clustering and PCA (**Fig. 1d, 1e and 1g**). Together, our time course phosphoproteomics data demonstrates that CCR2 activation rapidly stimulates numerous signaling events that collectively promote cell migration.

**Figure 2:**
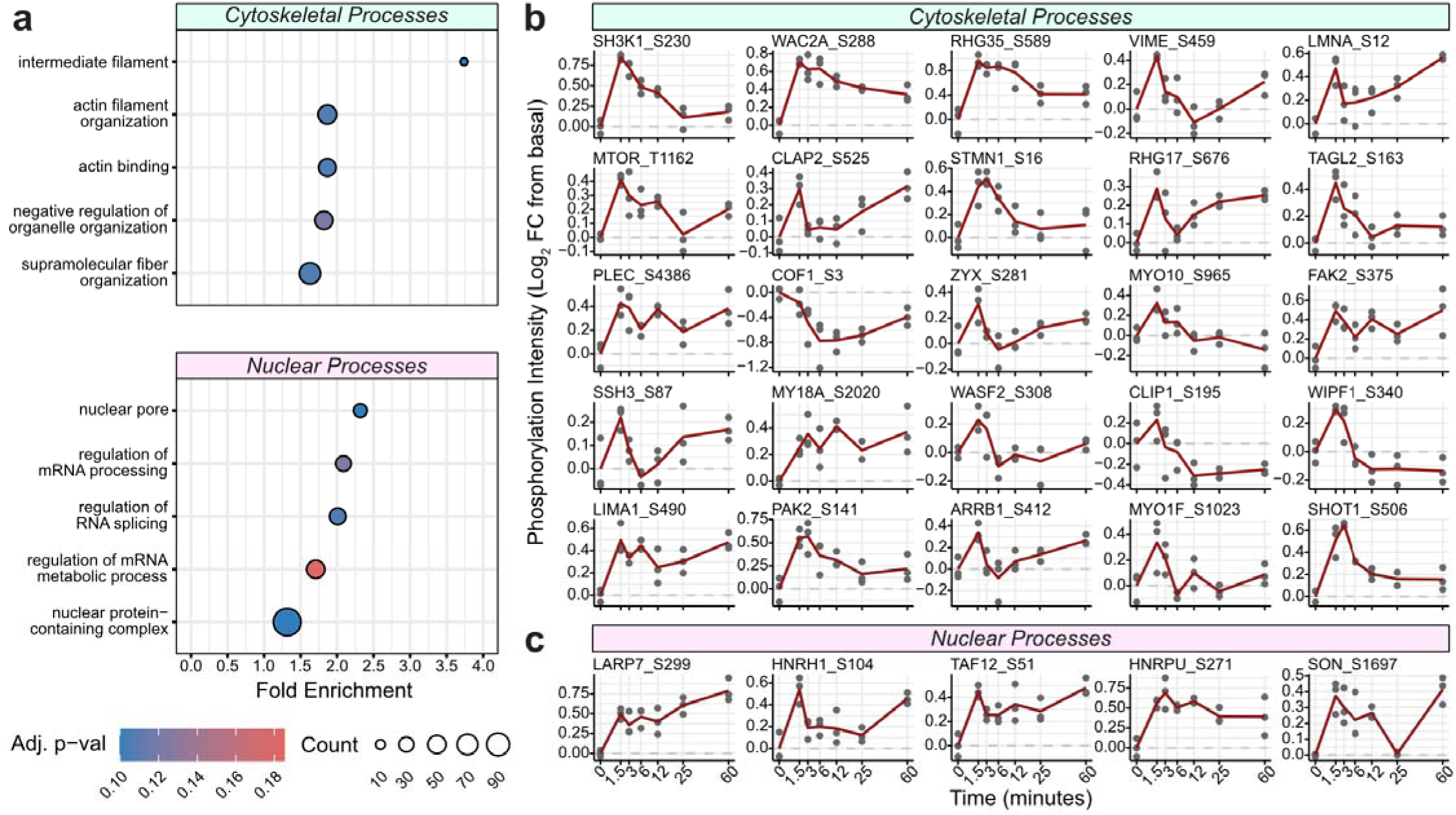
CCR2 activation initiates phosphorylation of actin-related and transcription-related proteins. **a**, Gene ontology analysis (molecular functions, cellular compartments, and biological processes) of the 398 ENTREZ-mappable phosphoproteins that were significantly regulated against the background of all 2,369 proteins detected in our dataset prior to filtering. Point positions indicate the linear fold enrichment compared to random; point size is relative to the number of proteins associated with each term; and color indicates Benjamini-Hochberg-corrected p-values. **b**,**c**, Representative phosphorylation time courses for phosphosites related to (**b**) cytoskeletal processes and (**c**) nuclear processes.

We also observed enrichment for some non-migratory cellular functions, most prominently involving proteins related to RNA processing, nuclear transport, and nuclear reorganization (**Fig. 2a** and **2c; Supplementary File 4**). Typically, these changes persisted for longer than those associated with cell migration, consistent with our understanding of the timescales of these biological pathways. These observations will be valuable for elucidating the mechanisms by which chemokines regulate protein expression, processing and localization^31–33^. Finally, we observed significant phosphorylation/dephosphorylation for numerous proteins that do not share any of these enriched functions. Although some of these are probably functionally inert in isolation, they may prime the cells for altered responses to other stimuli.

In one example of interplay between different stimuli, we observed CCL2-induced changes in phosphorylation of the C-terminal tail of CCR1 (**Supplementary Fig. 5a**), which regulates CCR1 internalization^34^. This suggests that CCR2 activation may alter the cellular response to CCR1 agonists through heterologous cross-desensitization. Indeed, in ERK phosphorylation experiments, stimulation of either CCR1 or CCR2 desensitized the other receptor to subsequent activation by its cognate agonist (**Supplementary Fig. 5b-d**). Notably, CCR1 desensitizes more readily than CCR2^35^, consistent with it being the dominant chemokine receptor in monocytes^36^.

### Network Modeling Reveals a Highly Divergent, Synchronized Signaling Cascade Downstream of Activated CCR2

Although time-resolved phosphoproteomics can reveal and functionally classify numerous phosphorylation events, it does not provide any direct information about the sequence and interdependence of the phosphorylation events, i.e., the signaling cascade. To generate a putative network resulting from CCR2 activation by CCL2, we used the set of observed phosphosites as input to a system-wide phosphorylation network modeling tool, PHONEMeS (PHOsphorylation NEtworks for Mass Spectrometry)^22,23^.

Briefly, PHONEMeS network modeling (**Fig. 3a**) involved: (1) building a prior knowledge network derived from the OmniPath and NetworKIN databases, supplemented with published predicted kinase-substrate relationships^37,38^ and a small set of user-defined connections not involving protein phosphorylation (e.g., activated G protein-dependent interactions) (**Supplementary File 5**); (2) refining the prior knowledge network to include only connections between proteins expressed in THP-1 cells (Human Protein Atlas^39^) and only the most direct pathways from CCR2 to each significantly-regulated phosphosite (the background network); and (3) iterative simulations to derive the optimal subset of the background network consistent with the full set of significantly-regulated phosphosites (the PHONEMeS network). Repeating the process for the phosphosites regulated at each successive time point enabled us to determine the time at which each connection first appeared in the network. Moreover, by performing 100 independent simulations, each with only 75% randomly selected significantly regulated and unregulated phosphosites, we estimated the necessity of each connection (represented as the consensus score; scale 1-100) in the resulting PHONEMeS network.

**Figure 3:**
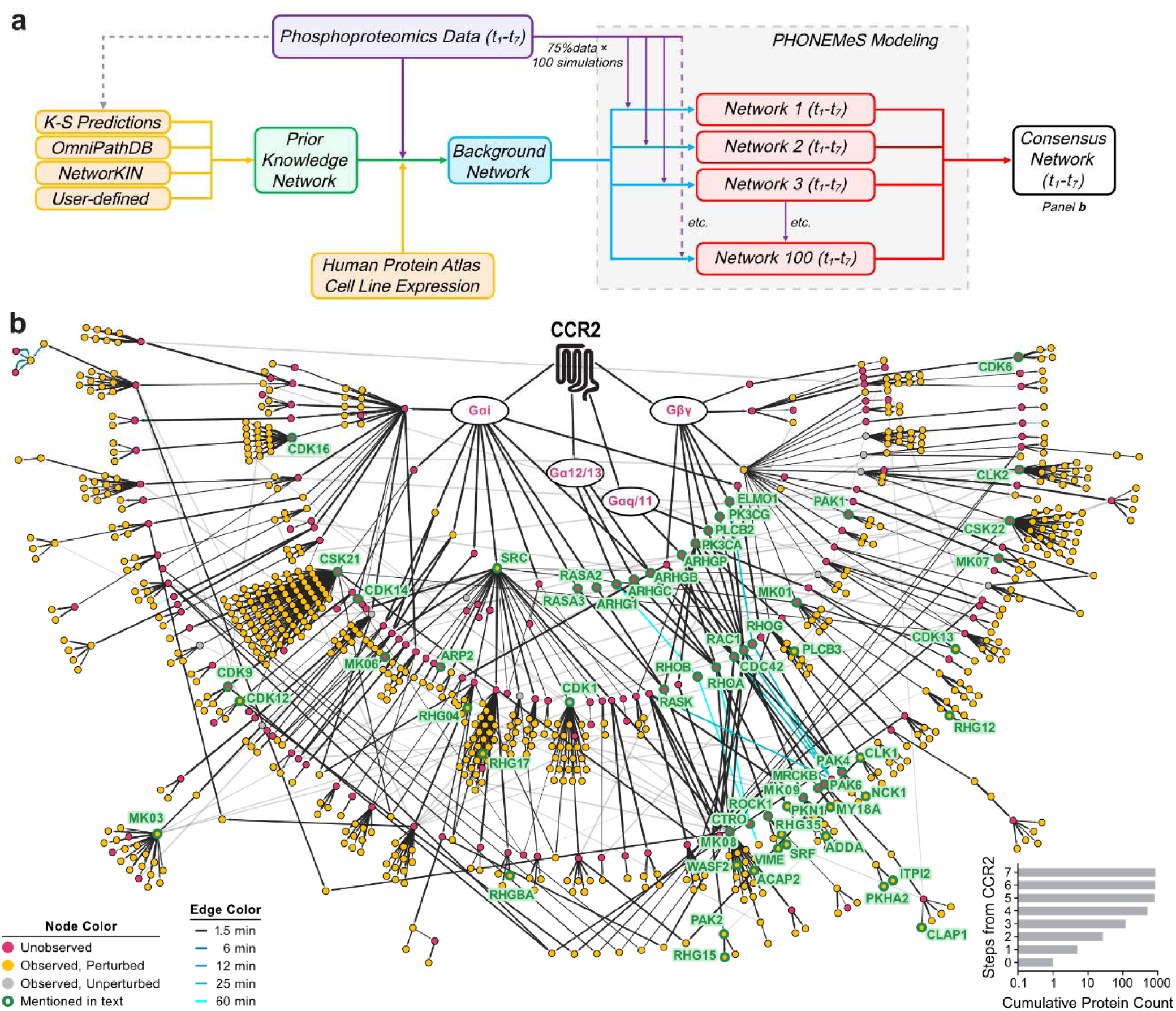
The CCL2-activated signaling network. **a**, Schematic of the PHONEMeS workflow for generation of a parsimonious signaling network. Significantly perturbed phosphosites observed through phosphoproteomics (purple) are incorporated through known interactions reported by multiple databases or kinase-substrate predictions (yellow). 100 networks (red) are generated through bootstrapping, each using 75% of significant phosphosites. K-S, kinase-substrate. **b**, The hierarchical representation of the resulting consensus network at the protein level. Nodes are proteins; edges represent known phosphorylation, dephosphorylation, or other regulatory events between them. Proteins harboring phosphosites that were significantly perturbed by CCL2 stimulation in THP-1 cells in this study are colored yellow. Proteins not observed in the phosphoproteomics data but identified by PHONEMeS as necessary links between CCR2 and perturbed phosphosites are colored rose. Proteins harboring phosphosites that were observed but not significantly perturbed in the present phosphoproteomics study are colored gray. Green circles around nodes and green labels signify proteins mentioned in the text. The time at which each protein-protein phosphorylation or regulatory event is first incorporated in the network is represented by increasingly vivid cyan colored edges. The inset (lower right corner) illustrates the exponential expansion of the network up to four steps from CCR2.

The complete PHONEMeS network (**Supplementary File 6)** consists of 1,630 phosphosites on 792 proteins, 72 non-phosphorylated proteins, and 2,876 interprotein connections. For presentation clarity, we focus on the consensus network, containing only the highest consensus score connections for each node, illustrated in **Fig. 3b** (protein level) and **Supplementary File 6** (protein and phosphosite levels). The consensus network incorporates pathways from CCR2 to 718 of the 1,387 significantly regulated phosphosites (and 594 of the 745 regulated phosphoproteins) in our dataset. Importantly, it also includes 111 proteins that were not observed in our data, most of them in the upstream part of the network (**Fig. 3b**). This illustrates the ability of the PHONEMeS approach to reveal biologically meaningful information not directly detected experimentally but required to explain the observed spectrum of protein phosphorylation.

The signaling network is highly divergent. As expected, the model indicates that CCR2 activates four families of G protein subunits: Gαi, Gα12/13, Gαq/11 and Gβγ. These G proteins activate 22 effector proteins (*vide infra*), which collectively activate 91 immediate downstream targets, illustrating the exponential expansion of the network (**Fig. 3b** inset).

Although the network includes proteins requiring up to seven sequential activation steps from CCR2, the vast majority (98%) of the 986 protein-protein connections appear in the network at the shortest time point measured (1.5 min), consistent with the above PCA and temporal clustering. Thus, the combined network structure and temporally resolved phosphorylation profiles indicate that CCR2-mediated signal transduction is essentially synchronized (on the time scale of <2 min), representing a rapid cascade of interconnected signaling events.

### G Protein Signaling is Divergent but Integrated

Receptor-activated heterotrimeric G proteins are known to pleiotropically couple to multiple downstream signaling pathways^40^. Our CCL2-dependent PHONEMeS network exemplifies this divergence of G protein signaling. The vast majority of the network is downstream of Gαi and/or Gβγ (**Fig. 4a**), consistent with the accepted view that chemokine receptor signaling is predominantly mediated by Gi-family proteins. Together, these subunits are seen to activate five main pathways (**Fig. 4b**). Gαi negatively regulates cAMP-dependent signaling and activates protein tyrosine kinases, most notably SRC^41^, which has 27 immediate downstream targets. Gβγ stimulates G protein receptor kinases (GRKs) and phosphoinositide signaling (PLC, PI3K). Both Gαi and Gβγ stimulate regulators of small GTPases of the Rho and Ras families. Although the Gαi and Gβγ pathways are largely distinct from each other, a subset of the network integrates signals from both, particularly at the MAP kinases MK01 (MAPK1 or ERK2), MK08 (JNK1) and MK09 (JNK2), each of which regulates at least ten experimentally observed significantly regulated phosphosites (**Fig. 4c**).

**Figure 4:**
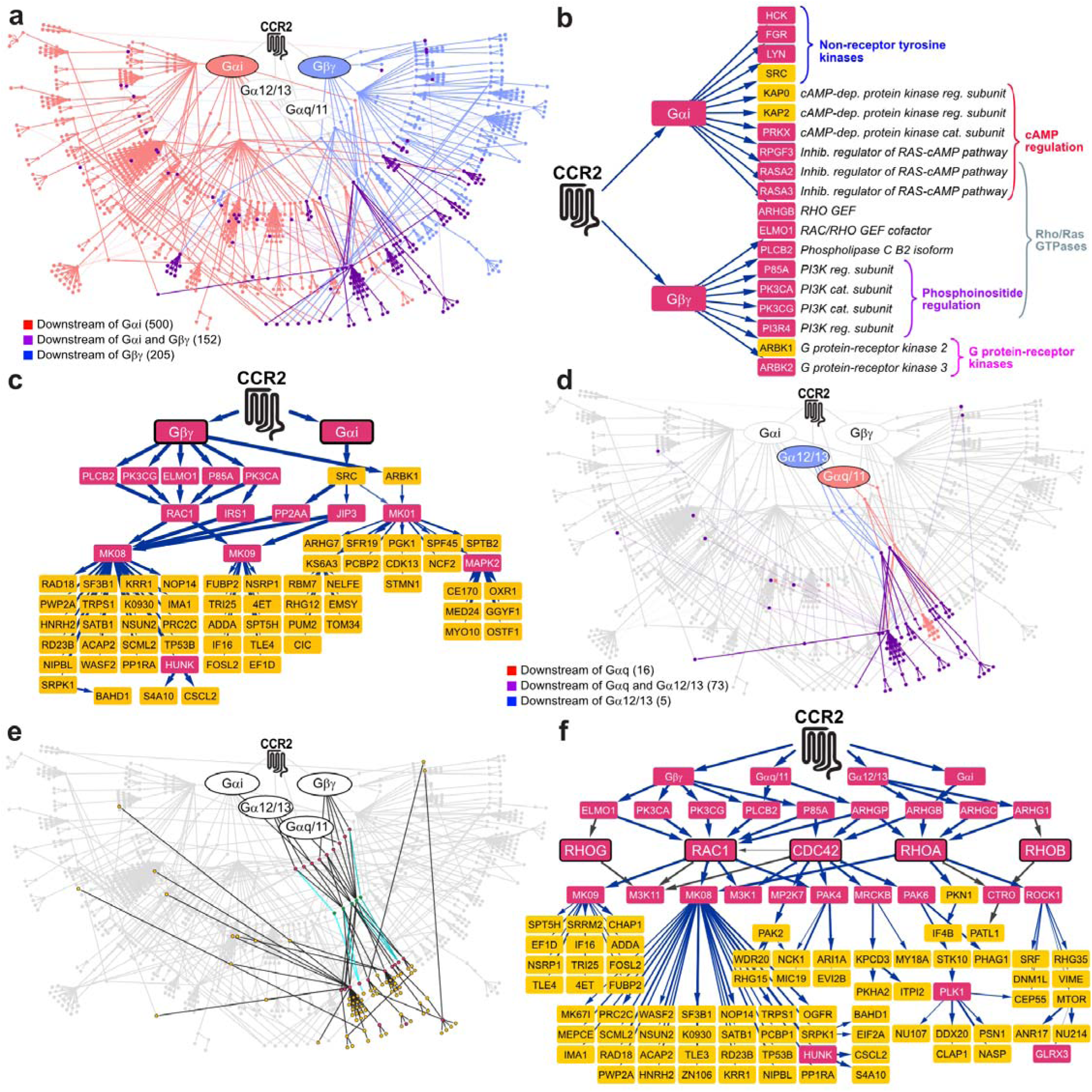
CCL2 stimulation regulates divergent chemotaxis-related proteins via G proteins and Rho GTPases. **a**, Network overview highlighting signaling pathways downstream of Gαi (red), Gβγ (blue) or both (purple). **b**, Five major protein families with distinct functions are located immediately downstream of Gαi and Gβγ. **c**, Subnetwork highlighting the Gαi and Gβγ signaling pathways leading to major MAPK signaling cascades. **d**, Network overview highlighting signaling pathways downstream of Gαq (red), Gα12/13 (blue) or both (purple). **e**, Network overview highlighting the Rho GTPase-mediated signaling pathways. **f**, Subnetwork illustrating phosphorylation-independent signaling from CCR2 to Rho GTPases and numerous regulated phosphoproteins downstream of Rho GTPases. For **b**, **c**, **e**, and **f**, node coloring is the same as in Fig. 3b.

In contrast to the Gαi/Gβγ-derived pathways, the Gα12/13 and Gαq/11-dependent signaling network appears more limited (**Fig. 4d**). There is substantial overlap between Gα12/13 and Gαq/11 signaling and further integration with Gαi/Gβγ in the early steps after G protein activation, converging on the Rho GTPases, as expected^42–44^. Thus, our network helps to define the mechanisms by which Gα12/13 and Gαq/11-dependent signaling regulates migration of leukocytes and other cell types^45^.

### Rho GTPases Are Central Chemokine Signaling Hubs

The PHONEMeS network includes five Rho family GTPases (RHOA, RHOB, RHOG, RAC1 and CDC42), signaling proteins with canonical roles in regulation of cell migration^15,46–51^. The connections from CCR2 to the Rho GTPases and from the Rho GTPases to 71 regulated phosphoproteins constitute a well-defined subnetwork (**Fig. 4e and 4f**). Signaling from CCR2-coupled heterotrimeric G proteins to RAC1, RHOA, RHOB, RHOG and CDC42 is mediated by nine effector proteins. Downstream of Gα12/13 and Gαq, these effectors include four Rho guanine nucleotide exchange factors, Rho GEFs [ARHGP/B/C/1]^1^ and the phospholipase C β2 isoform [PLCB2]. Downstream of Gβγ, the network features the ELMO1 (Engulfment and Cell Motility 1) adaptor for DOCK1 (dedicator of cyto-kinesis 1), a RAC1 and CDC42 GEF^52^, as well as PI3Kγ, which activates the Rho GTPases through PIP_3_-mediated GEF recruitment to the membrane^53^. In addition to the five Rho GTPases, the network includes the related small GTPase K-Ras [RASK], which has been identified as a central mediator of neutrophil polarization and migration^54,55^. Signal transmission from CCR2 to K-Ras occurs through RASA2 and RASA3, two Ras GAPs that act as direct effectors of Gαi^56^, consistent with the roles of these proteins in immune cell migration^54,55^.

The Rho GTPases collectively regulate 11 kinases that give rise to the observed protein phosphorylation in the downstream cascade. Among these kinases are well-established regulators of cell migration, including ROCK1, PAK4/6, citron kinase [CTRO], and MRCKB^15,57^. Notably, all connections upstream and immediately downstream of the Rho GTPases are present in 74-100% of simulated networks, indicating that these are the most likely pathways that account for the global phosphorylation patterns in our phosphoproteomics dataset. Thus, the Rho GTPase subnetwork demonstrates that phosphorylation-independent signaling upstream of the Rho GTPases regulates phosphorylation-dependent signaling downstream of the Rho GTPases.

Our network supports the critical involvement of Rho GTPases in the establishment and maintenance of cell polarity during migration. RAC1 and CDC42 are known to localize to the leading edge, promoting and coordinating actin polymerization and front extension, whereas RHOA activity is dominant at the trailing edge, stimulating actin-myosin contraction^57,58^. Consistent with this paradigm, RAC1 and/or CDC42 appear to regulate phosphorylation of WASF2, NCK1, ITPI2, MYO18 [MY18A], ACAP2, CLAP1, ADDA and PKHA2 (**Supplementary Table 1**). In addition, downstream of RHOA, the network includes phosphorylation of intermediate filament protein vimentin [VIME] and kinase PKN1, which negatively regulates intermediate filament assembly. We also observed phosphorylation of six Rho GTPase activating proteins [RHG04, 12, 15, 17, 35, and BA], which are expected to promote conversion of Rho GTPases to their inactive forms. These negative feedback mechanisms may help to coordinate cell polarization both spatially and temporally. While the kinase-substrate interactions for all of these proteins are derived from prior knowledge in existing databases, in many cases our data provide the first experimental evidence for the specific sites and/or time courses of phosphorylation and our network constitutes a set of testable hypotheses for their mechanisms of regulation.

### Divergent Pathways Regulate Actin Remodeling, Cell Polarization and Adhesion

The presence of Rho GTPase pathways important for leukocyte migration highlights the congruence of our PHONEMeS network with the current understanding of mechanisms regulating cell migration. However, these pathways only constitute a fraction of the network (∼12% of nodes; **Fig. 4e**), raising the question of whether they are sufficient to stimulate chemotaxis.

To identify additional signaling pathways that are regulated downstream of CCL2-activated CCR2, we performed semantic clustering of GO biological pathway terms for the proteins in the PHONEMeS network. This broadly reiterated the prevalence of pathways related to cell signaling, RNA regulation, and cytoskeletal processes (**Supplementary Fig. 6**). We then mapped proteins with GO terms related to cytoskeletal organization, adhesion and cell polarization onto the consensus network (**Fig. 5a-c**). For all three functional classifications, the relevant proteins are broadly spread across the divergent signaling network. Moreover, in some cases, multiple phosphorylation sites on the same chemotaxis-related protein are regulated by distinct signaling pathways, as shown for WASF2 (**Supplementary Fig. 7**). These observations suggest that the underlying cellular processes necessary for chemotaxis require activation of multiple divergent signaling pathways. This requirement for multiple pathways to be activated may enable tight regulation of cell migration. Moreover, it suggests that there are many potential opportunities for pharmacological intervention, as demonstrated below.

**Figure 5:**
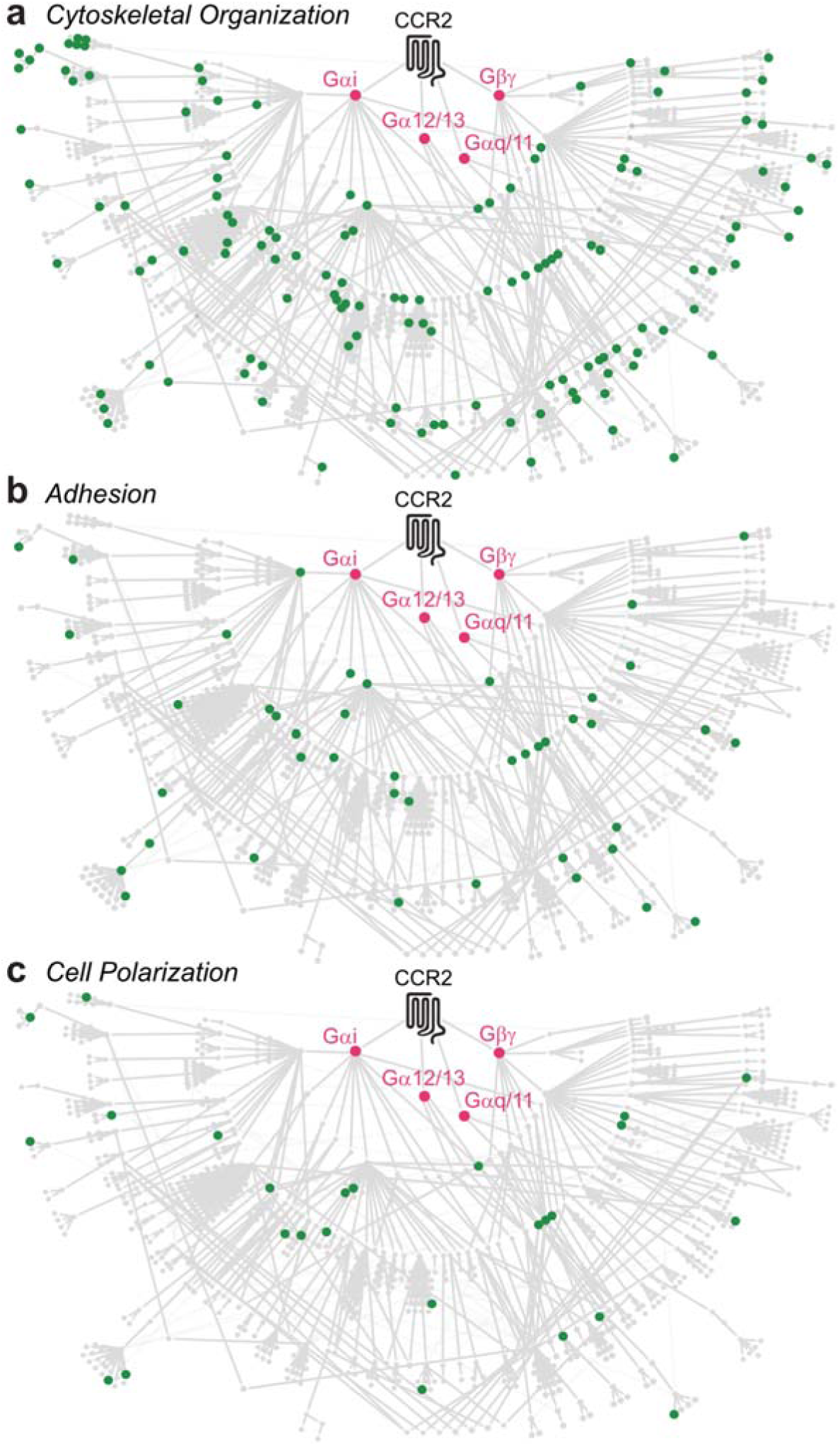
Proteins with functions related to cell migration are dispersed throughout the CCR2 signaling network. **a**,**b**,**c** The CCR2 signaling network in THP-1 monocytes is displayed with gray nodes (proteins) connected by gray edges. Signals originate from CCR2 (black) and are amplified via G proteins (rose). Green nodes indicate proteins related by Gene Ontology to (**a**) cytoskeletal organization, (**b**) adhesion, or (**c**) cell polarization.

### Blockade of Divergent Pathways Suppresses Cell Migration

We next investigated whether divergent branches of the signaling network are necessary for chemokine-stimulated THP-1 cell migration. We selected 35 kinases that could be targeted with 38 small-molecule inhibitors (**Fig. 6a; Supplementary File 7**), based on: (a) their broad distribution across the network; (b) their “betweenness centrality”, a metric of how frequently that protein directly links CCR2 to other proteins in the network; (c) their relationships to proteins with known (GO-annotated) functions in cell migration; and (d) the availability of validated, selective inhibitors. The target kinases are distributed across the network, together regulating 301 downstream proteins, more than a third of the network (**Fig. 6b**).

**Figure 6:**
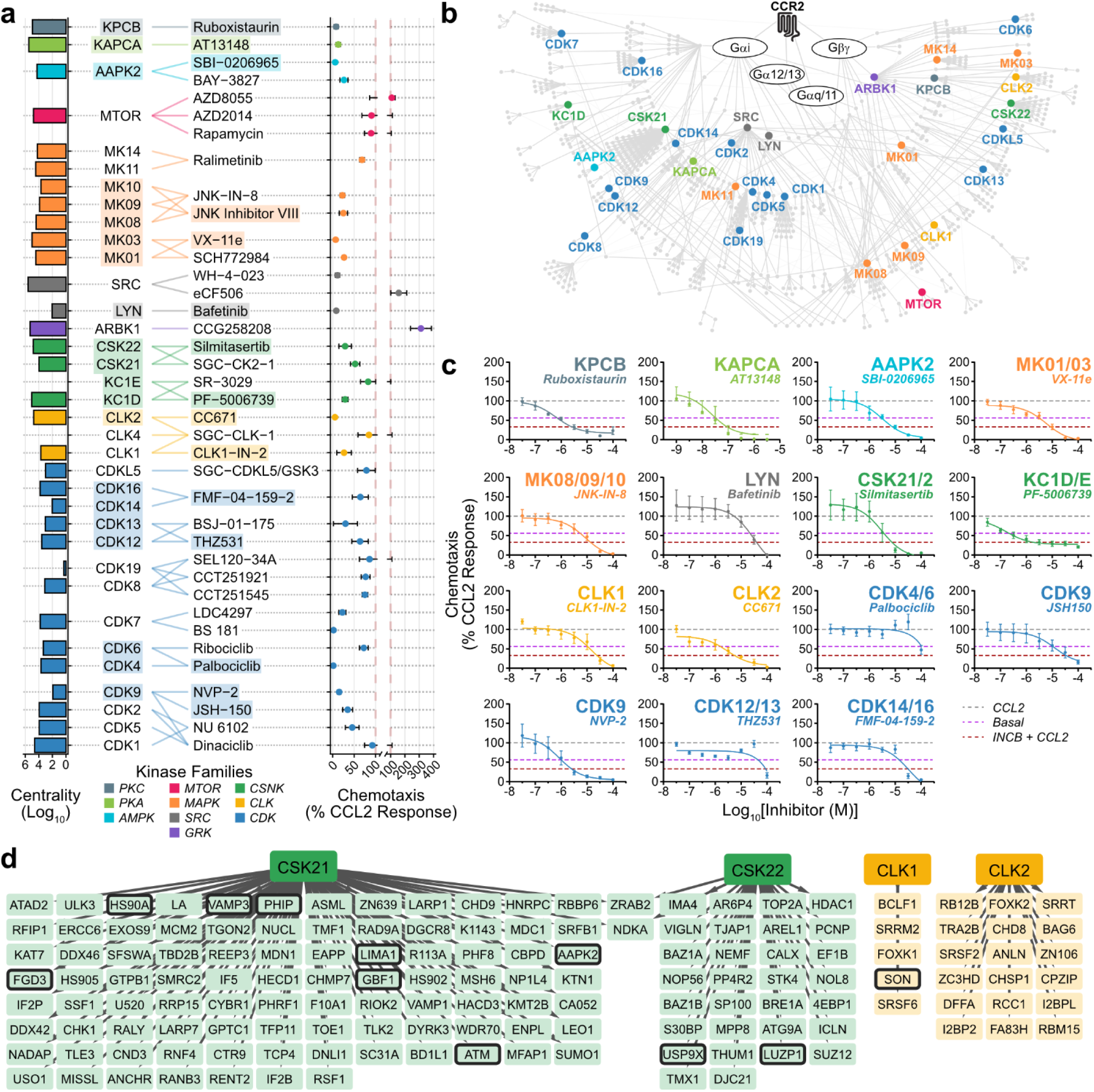
Kinase inhibitors targeting divergent branches of the CCR2 signaling network can successfully inhibit CCL2-driven chemotaxis. a, 38 kinase inhibitors targeting 35 kinases were selected for screening of inhibition of CCL2-stimulated chemotaxis (right) on the basis of their betweenness centrality (left). Points and error bars represent mean and SEM of n=3-4 experiments, while the red dashed line represents the CCL2 response. Points and centrality bars are colored according to the kinase family. Shaded targets and inhibitors indicate those selected for further validation. b, Selected kinases are distributed widely throughout the signaling network, and represent strong centers of signaling. Nodes are colored according to the kinase family (as in a). c, 15 inhibitors targeting 22 kinases (or kinase sub-families) display concentration-dependent inhibition of CCL2-stimulated THP-1 monocyte chemotaxis. Points and error bars represent mean and SEM of n=4 inhibitor experiments and n=14-16 for controls. Dashed lines: gray, CCL2 response; magenta, basal response; red, CCL2 response after 2 h pretreatment with 10 μM INCB. d, The downstream targets of CSK21, CSK22, CLK1, and CLK2 are shown. Nodes are colored according to the protein family, as in a. d, The downstream targets of CSK21, CSK22, CLK1, and CLK2 are shown. Inhibitor target kinases are colored according to the protein family, as in a, with paler tones indicating substrates. Black borders indicate substrates involved (based on GO terms) in cytoskeletal organization, adhesion, or cell polarization.

We screened all 38 kinase inhibitors in a CCL2-stimulated chemotaxis assay (**Fig. 6a**; **Supplementary Fig. 8**). We then evaluated 15 kinase inhibitors in concentration-response curves (**Fig. 6c**), chosen for their strong inhibition of THP-1 chemotaxis, negligible effects on cell viability, and high compound solubility. Among these, 13 inhibitors, targeting 12 kinases (or kinase subfamilies), suppressed CCL2-stimulated migration by more than 50%. These targets included CSK21/22, MK03/01 (ERK1/2), CDK9 and CLK1/2, which were also highlighted by the Kinase Enrichment Analysis of our dataset (**Supplementary Fig. 3**). CDK4/6 and CDK12/13 inhibitors did not suppress chemotaxis in a concentration-dependent manner (**Fig. 6c**). These chemotaxis inhibition data are robust, also confirming previous data linking protein kinase A, protein kinase C, MK08/09 (JNK1/2) and LYN with leukocyte migration^59–63^. The inhibition of ARBK1 (GRK2) *increased* chemotaxis rather than decreasing it (**Fig. 6A**). This is consistent with the role of GRK2 in CCR2 phosphorylation and desensitization^64^, and likely occurred via inhibition of these processes, resulting in increased CCR2 presence at the plasma membrane and sensitivity to CCL2.

Notably, most of the kinase inhibitors reduced migration to well below basal levels. Our inhibition studies (**Fig. 1a**), as well as prior studies^65^, had indicated that basal migration (in the absence of any ligand) is largely dependent on CCR1, suggesting that the targets of these inhibitors play roles in CCR1- as well as CCR2-mediated monocyte motility.

In summary, our kinase inhibition data confirm that CCR2-mediated migration requires phosphorylation-dependent signaling via multiple divergent pathways, under the control of both recognized and novel kinases. Our network will guide future identification of the relevant substrates of these kinases and the detailed mechanisms by which they contribute to chemokine-driven monocyte migration.

## Discussion

Despite the key roles that chemokine receptors play in guiding immune cells and fueling inflammatory cascades, their therapeutic targeting has so far had limited success^15,66,67^. Among the commonly cited reasons for clinical failures are incomplete inhibition of migration due to the redundancy of the chemokine system, insufficient occupancy of receptors by drugs, and the difficulty translating from rodent models to humans due to different roles of chemokine-receptor pairs across species^15,67–69^. In these circumstances, targeting downstream proteins that transduce the signals from activated chemokine receptors may provide a more efficient approach for treating inflammation. However, our understanding of chemokine-receptor-dependent migratory signaling in immune cells is currently insufficient to identify suitable targets for this blockade strategy.

To elucidate the intracellular signaling pathways downstream of CCR2, and to assist the discovery of novel targets for autoimmune and inflammatory diseases, we performed an unbiased time-resolved phosphoproteomics study. Moreover, considering that it is not possible to capture all intracellular signaling events with phosphoproteomics, we interpreted the changes in protein phosphorylation by computational network modeling, in the context of known signaling protein relationships. This revealed an extensive, divergent signaling network resulting from CCR2 activation in monocytes.

Our network recapitulates the key features of the currently prevailing theory of chemotactic signaling, including the dominant involvement of Gαi (cAMP-inhibitory) and Gβγ proteins, and the central roles of Rho GTPases. Recent studies suggest that the constituent processes that define chemotaxis - such as directional sensing and actin remodeling with membrane protrusion formation - can be functionally decoupled from chemical stimulus-induced, non-directional cell motility (i.e., chemokinesis)^70^. For example, while chemotaxis requires the dissociation of Gβγ subunits from Gαi proteins^71^, CCL2-induced increases in actin remodeling are Gi-independent^72^. Similarly, in epithelial-derived cancer (HeLa) cells, the directionality of CXCL12-induced migration, but not non-directional chemokinesis, is controlled by the mTORC1 and mTORC2 complexes^73^. Moreover, while chemotaxis and chemokinesis can occur downstream of non-G protein-coupled receptors (e.g., EGFR) without Gi involvement^74^, in all cases the signals from surface receptors are transmitted to the actin assembly via the small GTPases including RAC, RAC, RHO, and CDC42^46,75^. Our network recapitulates the activation of these GTPases as a common outcome that integrates diverse receptor-initiated signals and redirects them towards a critical subset of the pathways necessary for cytoskeletal rearrangements and cell motility.

By filling the gaps in the proteomics data, the PHONEMeS network enabled prospective predictions regarding the roles of kinases that were not directly observed in phosphoproteomics. While some were expected (e.g., MK03/01, or KAPCA, the catalytic subunit of PKA), others have not been previously reported to be involved in chemotaxis. For example, our signaling network included multiple members of the cyclin-dependent kinase (CDK) family. CDKs are well-known regulators of the cell cycle and proliferation, and have been previously implicated in cancer cell migration and invasion^76–78^ but their role as leukocyte migration factors is less recognized^63,79,80^. Experiments with CDK inhibitors confirmed the role of CDK9 and CDK14/16 in transmitting the chemotactic signals downstream of CCR2. Similarly, our inhibition experiments revealed roles of casein kinases 2A and 2B (CSK21 and CSK22) and CDC-like kinases 1 and 2 (CLK1 and CLK2), which have numerous direct substrates in our PHONEMeS network (**Fig. 6d**). Casein kinases are broad specificity Ser/Thr kinases, whereas CLK1 and CLK2 are members of the dual-specificity (Tyr and Ser/Thr) kinase family. Both casein kinases and CDC-like kinases have been previously implicated in migration and tissue-invasion of cancer cells^81–84^ but they are not known to regulate leukocyte migration. In our experiments, inhibition of all four kinases robustly suppressed CCL2-induced chemotaxis of THP-1 cells.

Our inhibition data clearly demonstrate that individual blockade of various kinases spread across the signaling network can suppress chemotaxis. Thus, the activity of each successful target is necessary but not sufficient for chemokine-stimulated cell migration. These results support the paradigm in which chemotaxis emerges as an integrated cellular response to divergent signaling pathways. Considering that our study focused on CCR2-mediated signaling in monocytes, it will be important to investigate whether the same paradigm applies to signaling via different chemokine receptors in other types of leukocytes.

Arguably the most important implication of this paradigm is that effective blockade of chemokine-driven inflammation might be achievable by inhibiting any one of numerous different proteins within the divergent network. Indeed, considering that several inhibitors suppressed both CCR2-mediated chemotaxis and CCR1-mediated basal migration, our data suggests that blockade of key signaling proteins will be more effective than the traditional approach of targeting individual receptors. Furthermore, the methodology used in this study could be readily applied to different leukocyte types to identify mechanisms that regulate chemotaxis in specific cell types, thus creating opportunities for cell type-specific interventions.

In summary, our chemokine-driven signaling network reveals fundamental insights into the biochemical coordination of leukocyte migration and provides an initial “road map” for identification of suitable targets for anti-inflammatory drug discovery.

## Methods

### Chemicals and biochemicals

CCL2 and CCL5 were purchased from PeproTech (cat. nos. 300-04 and 300-06) or ChemoTactics (cat. nos. CCL2-100ug and CCL5-100ug).

Reagents used for cell culture and cell-based assays were: RPMI 1640 Glutamax (Gibco, cat. no. 61870-127); RPMI 1640 medium, no phenol red, no glutamine (Gibco, cat. no. 32404014); South American Fetal Bovine Serum (FBS), gamma-irradiated (Gibco cat. no. 10499-044, lot 1946455K); sodium pyruvate (Gibco cat. no. 11360070); glucose (Gibco cat. no. A2494001); Glutamax (Gibco Cat, #61870-127); HEPES (Gibco, cat. no. 15630080); Hank’s Balanced Salt Solution (HBSS) (Thermo Fisher, cat. no. 14175095); penicillin/streptomycin (P/S) (Sigma cat. no. 70002278); magnesium chloride hexahydrate (Sigma-Aldrich, cat. no. M2670-100G); bovine serum albumin (BSA) (Sigma Aldrich, cat. no. A7906-100G); and CellTiter-Glo 2.0 Assay (Promega cat. no. G9242).

Reagents used in sample preparation for mass spectrometry were: Tris-buffered saline (TBS) (ThermoFisher Scientific, 28358); sodium deoxycholate (Sigma, D6750), methanol (Merck, 1.06007); 1M HEPES (Gibco, cat. no. 15630080) at pH 8.5; 500 mM Bond-Breaker TCEP (ThermoFisher Scientific, 77720); 500 mM DTT (Astral, C-1029-5g); chloroacetamide in powder (Sigma, C0267-100G); Trypsin Gold (Promega, V528X); LysC (Wako Chemicals, 129-02541); formic acid (Sigma, 56302-50ML-F); acetonitrile (ACN) (ThermoFisher Scientific, Optima LC/MS, A955-4); bicinchoninic acid (BCA) kit (ThermoFisher Scientific, #23225); isopropanol (ISO) (Fluka, V800228); potassium dihydrogen phosphate (Sigma, 1048731000); trifluoroacetic acid (TFA) (Sigma, T6508); ammonium hydroxide solution (Sigma, 338818-100ML); titanium dioxide (TiO_2_) beads (Titansphere Phos-TiO Bulk 10 μm; GL Sciences, 5010-21315).

Chemokine receptor antagonists INCB3344 (cat.no. A11465) and BX471 (cat.no. A14409) were purchased from Sapphire Biosciences. Kinase inhibitor suppliers and catalogue numbers are listed in **Supplementary File 7**.

### Cell lines and cell culture

THP-1 monocytes were purchased from ATCC (product number TIB-202) and used for all experiments. THP-1 cells were maintained in culture medium (RPMI 1640 Glutamax, supplemented with 10% heat inactivated FBS, 1% P/S, 1 mM sodium pyruvate, 25 mM glucose) at 37°C and 5% CO_2_ and split every 2-4 days. To ensure optimal growth, a density between 0.2 and 1 million cells/mL was maintained.

### qRT-PCR analysis of chemokine receptor expression

RNA extraction was performed using the RNeasy Plus Mini Kit (Qiagen, cat. no. 74134) according to the manufacturers’ protocol. cDNA was generated using the High-Capacity cDNA RT Kit (Applied Biosystems/Life Technologies, cat. no. 4374966) following the instructions of the manufacturer. To design the primers for qRT-PCR (**Supplementary Table 2**), Harvard Primer Bank or NCBI Primer-BLAST was used. Reactions were performed in 10 μL containing 5 ng cDNA and 0.5 μM primers using PowerUp SYBR Green Master Mix (Thermo Fisher, cat. no. A25741). The reaction was carried out in a Hard-Shell 96-well PCR Plate (Bio-Rad, cat. no. HSP9601) with the Bio-Rad CFX 96 real-time PCR detection system. 18S was used as internal control. Relative gene expression was quantified using the established 2^−ΔΔCT^ method^85^, compared to CCR1 expression.

### Chemotaxis and inhibition of chemotaxis

Chemotaxis assays were carried out using Merck 96-well Transwell plates with a pore size of 5 µm (cat. no. MAMIC5S10). The lower chambers were loaded with chemokines in RPMI buffer at the indicated concentrations (or buffer alone) and the upper chambers were loaded with 400,000 THP-1 cells in RPMI buffer. The plates were placed in a humidified container during incubation at 37 °C with 5% CO_2_ for 2 hours. CellTiter-Glo 2.0 was used to quantify the cells that had migrated into the bottom chamber. Luminescence was measured using a Pherastar plate reader (BMG Labtech) equipped with the luminescence module (wavelength 560-600 nm).

To validate CCR2- or CCR1-specific activity, cells were preincubated with INCB3344 (1 μM) or BX471 (1 μM), respectively, corresponding to 100-fold above their IC_50_ values, for 20 min prior to treatment with the chemokine (CCL2 or CCL5, respectively). The chemotaxis response is expressed as the mean chemotactic index, calculated as fold-change over basal.

To evaluate the effects of small-molecule kinase inhibitors on CCL2-stimulated chemotaxis, the cells were incubated with the inhibitor at the indicated concentrations (or vehicle) for 2 hours mins at 37°C, 5% CO_2_ before addition of CCL2 (10 nM), then incubation and measurement of migrated cells as described in the previous paragraph. Concentration-response data for small molecule inhibitors were adjusted for high concentration DMSO effects (up to 1%), normalized between 0% (no chemokine or inhibitor) and 100% (10 nM CCL2, no inhibitor), and fit to a logistic single-site regression model using GraphPad Prism 9.0.

### ERK1/2 phosphorylation assay

THP-1 cells grown in culture media were harvested and washed using the assay buffer (HBSS supplemented with 0.1% BSA, 1mM CaCl_2_ and 1 mM MgCl_2_). The cells were added at 2 × 10^6^ cells/mL (100,000 cells per well) to a 96-well clear culture plate (Nunc, ThermoFisher Scientific, cat. no. NUN167008) and serum starved at 37°C with 5% CO_2_ for 2 hours. The cells were then treated with chemokine at the indicated concentrations (or buffer alone) for 5 min at 37 °C. The reaction was terminated by adding 25 μl of SureFire lysis buffer per well and shaking at ∼400 rpm for 10 min. Detection of ERK1/2 phosphorylation in cell lysates was carried out using the AlphaLISA SureFire Ultra phosphoERK1/2 assay kit (Perkin Elmer, cat. no. ALSU-PERK-A500) according to the manufacturer’s instructions. The fluorescence signals were measured using a Pherastar plate reader (BMG Labtech) with standard AlphaLISA settings.

To validate CCR2- or CCR1-specific activity, cells were preincubated with INCB3344 (1 μM) or BX471 (1 μM), as for chemotaxis assays described above. Data were normalized between 0% (no treatment) and 100% (100 nM CCL2).

Interplay between CCR1 and CCR2 was assessed by measuring ERK1/2 phosphorylation after sequential treatments with agonists of each receptor. Briefly, THP-1 monocytes were plated in 6-well plates and serum-starved for 4 h. Cells were then subjected to a primary stimulation with CCL2 or CCL5 (100 nM), or buffer control, for 10 min, 5 min, or 0 min. Cells were then centrifuged, washed, and resuspended in fresh medium before plating at 2 × 10^6^ cells/mL in a 96-well plate. Exactly 8 min after the variable primary stimulation period, secondary stimulation was performed using CCL2 or CCL5 (100 nM), or buffer control, for 5 min, resulting in total intervals of 18, 13 or 8 min between primary and secondary stimulations. pERK levels were quantified as described above.

### Phosphoproteomics sample preparation

THP-1 monocytes were cultured in RPMI with 10% heat-inactivated FBS. Two million cells resuspended in serum-free media were seeded per well in two wells of a 6-well plate and serum-starved for 4 h before being stimulated with 100 nM CCL2 for 1.5, 3, 6, 12, 25 or 60 min or treated with vehicle (basal control). For INCB3344 treatments, cells were first pretreated with 1 μM INCB3344 for 20 min prior to CCL2 stimulation. Cells were then washed with ice-cold TBS and lysed on ice in a solution comprising 4% sodium deoxycholate and 100 mM Tris (pH 8.0), employing a probe sonicator (Soniprep 150, MSE; amplitude 10) for three cycles of 30 s. Subsequently, the protein concentration was quantified using a BCA assay kit according to the user manual. Three independent biological replicates were performed.

For protein digestion, 300 µg of protein was subjected to denaturation and alkylation using 10 mM tris(2-carboxyethyl) phosphine and 40 mM chloroacetamide at 95°C for 5 min. The lysate pH was adjusted to 8.0, and trypsin (Trypsin Gold, Promega) was introduced at a ratio of 1:100 (trypsin: protein, w/w), followed by incubation at 37 °C with agitation on a Thermomixer (Eppendorf) for 16 hours. Peptide fractionation was performed using a ZORBAX 300 Extend-C18 column (4.6 x 250 mm, 5 µm or 4.6 x 50 mm, 3.5 µm) connected to an HPLC 1100 (Agilent Technologies). 66 peptide-containing fractions were collected between 7 and 72 min and pooled non-contiguously, as previously described^86^, resulting in a total of 12 fractions. The samples were subjected to phosphopeptide enrichment using TiO_2_, as described previously^87,88^, and reconstituted in mass spectrometry sample resuspension buffer.

### Proteomics data collection

Liquid chromatography-mass spectrometry/mass spectrometry (LC-MS/MS), in data-dependent acquisition (DDA) mode, was used to quantify enriched phosphopeptides. Briefly, peptides from enriched phosphopeptide samples were separated using a Dionex UltiMate 300 RSLCnano system using an Acclaim PepMap RSLC analytical column (75 µm x 50 cm, nanoViper, C18, 2 µm, 100Å; Thermo Scientific) coupled with an Acclaim PepMap 100 trap column (100 µm x 2 cm, nanoViper, C18, 5 µm, 100Å; Thermo Scientific). The LC gradient, comprised of 0.1% formic acid (buffer A) and 80% ACN 0.1% formic acid (buffer B), was linearly ramped, from 7% to 37.5% buffer B over 120 min, prior to nebulization and electrospray ionization using a nano electrospray source (ThermoFisher Scientific) with a distal-coated fused silica emitter (New Objective) set to 1.7 kV capillary voltage. Ionized samples were analysed using a Q-Exactive HF hybrid quadrupole-Orbitrap mass spectrometer (Thermo Scientific, Bremen, Germany). The spectrometer operated in DDA mode with 12 fragmentation spectra per duty cycle. Precursor scans were conducted with a resolution of 120,000 (at 200 m/z) within the range of 375 to 1,575 m/z, using an ion target of 3 × 10^6^ and a maximum injection time of 54 ms. Fragmentation spectra were obtained with a normalized collision energy of 27 V at a resolution of 30,000, employing an isolation window of 1.4 m/z, a starting mass of 120 m/z, and acquiring only charge states 2-5. A dynamic exclusion of 15 s was applied for all acquired precursors.

### Phosphoproteomics data processing

MaxQuant (v1.5.2.8) and its search engine Andromeda was used to deconvolute MS/MS spectra into candidate peptides. Briefly, DDA files were searched against the September 2019 release of the UniProt reference human proteome. Cysteine carbamidomethylation was designated as a fixed modification, while methionine oxidation, N-terminal acetylation, and serine/threonine/tyrosine phosphorylation were defined as variable modifications. The search allowed for up to two missed cleavages and a mass tolerance of 20 ppm for the initial search, resulting in 12,282 phosphosites from 2,577 proteins with an FDR < 0.01.

For downstream analyses, only phosphosites with localization probabilities ≥75% were considered. Spectral intensities (corresponding to abundance of phosphorylated species) were transformed into log_2_ scale and median-centered (**Supplementary Fig. 9a-b**). Samples collected on different days (each involving a single set of stimulation and inhibition conditions) were treated as technical batches; technical variation between these batches (“batch effects”) was removed using linear modeling as implemented in the *limma v3.62.2* R library^89,90^. Quality of batch effect removal was controlled by comparing adjusted Rand indices^91^ between the experimental variables and the sample-to-sample correlation dendrogram cut at different heights (**Supplementary Fig. 9c-d**)^92^. Batch effects could not be removed for 1,722 phosphopeptides due to missing data, resulting in a normalized dataset of 8,079 phosphopeptides. Unambigious phosphopeptides were mapped to phosphosites. Subsequent filtering for phosphosites that had measurements in at least 2 out of 3 replicates across all time points, excluding INCB pretreated samples, and the most significant multiplicity per phosphosite reduced the set to 6,297 phosphosites from 2,115 proteins.

### Phosphoproteomics data analysis

For each phosphosite in the filtered dataset, statistical significance of its variation across all chemokine/time conditions, excluding the INCB3344 treated condition, was assessed using moderated ANOVA with Benjamini-Hochberg adjustment for multiple comparisons, as implemented in *limma v3.62.2*. This led to the identification of 630 significantly varied phosphosites (FDR < 0.05).

For **Fig. 1d**, vectors of log_2_ fold-changes of intensities in the treated conditions, relative to the basal, were compared to each other using a Euclidean distance metric, hierarchically clustered using McQuitty linkage (*hclust()* function in R), and plotted as a heatmap using the *pheatmap v1.0.13* library in R, with optimal leaf ordering (OLO) as implemented in the *dendextend::seriate_dendrogram()* function.

For time course clustering (**Fig. 1e** and **Supplementary Fig. 4)**, pairwise similarity between the vectors of log_2_ intensities for the 3,096 phosphosites detected at all time points (**Fig. 1d**) was calculated using the extra sum-of-squares F test^93^. F test p-values were transformed to negative log_10_ scale (so that larger numbers corresponded to more distinct profiles) and used as phosphosite-phosphosite profile distances. The resulting distance matrix was used to hierarchically cluster phosphosites with complete linkages (*hclust()* function in R) according to their variation profiles across all chemokine-treated conditions. A cutoff height of 3.5 was used to separate proteins into 20 clusters.

For **Fig. 1f**, the matrix of log_2_ intensities of the 630 significantly varied phosphosites (21 intensities each) was subjected to principal component analysis (PCA) as implemented in *pcaMethods v1.98*^94^ with missing values estimated by *SVDimpute*^95^. The first two PCs for each sample were plotted against each other to provide a high-level overview of consistent variation patterns across samples.

For **Fig. 2a**, over-representation of Gene Ontology Biological Process (BP), Cellular Compartment (CC), and Molecular Function (MF) terms was analyzed using the *clusterProfiler v4.14.6* R library^30^: the 400 significantly varied phosphoproteins were mapped to gene-based ENTREZ IDs and annotated against the background of all 2,369 proteins reliably detected in our dataset. Significances of overrepresented terms were adjusted using Benjamini-Hochberg correction for multiple comparisons.

### PHONEMeS network generation

A linear model was fit to all time points using *limma v3.62.2*^89,90^. A contrast matrix was then constructed for pairwise comparisons between the basal state and each of the follow-up time points, and used to compute the statistical significance of each phosphosite intensity at each time point relative to basal / untreated state using empirical Bayes moderated t-statistics with Benjamini-Hochberg adjustment. The p-values were normalized relative to 0.05 to produce scores used for integer linear programming (ILP) optimization in PHONEMeS-ILP^96^ (negative for sites significantly regulated, relative to basal, at the given time point, and positive otherwise).

A “prior knowledge” network was compiled from known kinase-substrate, phosphatase-substrate, and other regulatory relationships reported by *Omnipath*^97,98^ and *NetworKIN*^99^. For kinase- and phosphatase- mediated relationships, the nature of the enzymes was confirmed based on *UniProtKB* annotations. The network was complemented by predicted kinase-substrate pairings generated from the phosphopeptides presented in the current study and two kinome-scale aggregated mass spectrometry datasets^37,38^. Briefly, phosphosites (and their parent phosphopeptides) from all three studies were scored against all possible serine/threonine/tyrosine kinases using the *kinase-library v1.1.0* Python package. To account for the varying degrees of phosphorylation ubiquity for each phosphosite, these scores were converted into z-scores for each individual phosphosite. A kinase-phosphosite pairing (K-S) was added to the PKN if the kinase z-score, *Z_k_*(*S*), satisfied the following criteria: *Z_k_*(*S*) ≥ *Min*(*max*(*Z_i_*(*S*)), 3.1)-0.1, where the maximum is taken over all kinases (*i*) in relation to phosphosite *S*. This filter ensured that phosphosites that were poorly phosphorylated across the kinome were linked to only few kinases, whilst phosphosites that were ubiquitously phosphorylated were linked to many high-scoring kinases (note: z-score above 3 corresponds to the top 0.15% of all kinases). Furthermore, the “prior knowledge” network was filtered to only include proteins with detectable mRNA expression in the THP-1 cell line, as reported by the *Human Protein Atlas*^100^. Relationships involving CCR2, heterotrimeric G protein subunits, and selected relationships involving small GTPases were manually curated into the “prior knowledge” network **(Supplementary File 5)**.

To define the search space for PHONEMeS-ILP, a “background” network was subsetted from the “prior knowledge” network. Edges comprising all simple / acyclic paths of length of 7 or shorter and connecting CCR2 to experimentally observed phosphosites were included in this “background” network. Notes without incoming edges were removed.

The *PHONEMeS-ILP*^96^ tool was run using IBM *CPLEX* to connect the experimentally observed and significantly regulated phosphosites to CCR2, using edges from the “background” network. The PHONEMeS-ILP objective function is designed to minimize the weighted sum of nodes and interactions, where the inclusion of observed significantly regulated phosphosites is rewarded proportionally to their normalized p-values as above, the inclusion of observed non-regulated sites is penalized, and there is an additional penalty for including unnecessary edges. Phosphosites that were significantly regulated (p<0.05) at the earliest experimental timepoint (1.5 min) were incorporated via the first iteration, and those significantly regulated at later time points were sequentially integrated into the assembled network until all time points had been incorporated. This sequential network construction was performed 100 times, each time using 75% of the significantly perturbed phosphosites, selected at random, to build 100 partial networks. A consensus network was compiled by merging all nodes and edges in the partial networks, with the number of partial networks in which each specific edge has occurred interpreted as this edge’s “consensus score”. The full network (**Supplementary File 6**) contained 2,494 nodes (1,630 sites and 864 proteins) and 5,141 edges. By the design of the PHONEMeS network, its nodes (proteins and phosphosites) are classified as observed-and-significantly-regulated in the phosphoproteomics experiment (1,037 phosphosites in our network), observed-and-not-regulated (13 phosphosites in our network), and unobserved / added by the algorithm based on the prior knowledge network to ensure connectivity (762 nodes in our network of which 580 are phosphosites, 111 proteins with phosphosites, and 71 proteins without phosphosites).

### Generation of filtered consensus PHONEMeS network

The excessive number of low-consensus edges in the full PHONEMeS network necessitated filtering and simplification for the purposes of effective display and analysis (**Fig. 3b**). First, the network was cast from the level of phosphosites to the level of proteins. For this, implied nodes representing terminal phosphoproteins were added. Then, each phosphosite was replaced with a new edge connecting its upstream kinase or phosphatase and its parent (substrate) protein, retaining the consensus score of the phosphosite’s incoming edge. In cases where a single upstream kinase/phosphatase was connected to a single substrate protein via multiple phosphosites, the edge was assigned the maximum consensus score from the earliest time points. Phosphoproteins were considered observed-and-regulated if they harbored at least one observed-and-regulated phosphosite.

A simplified “jittered arborescence” network was generated by selecting the highest-consensus incoming edge for every protein node (the Chu-Liu/Edmonds algorithm^101,102^, implemented in R). To account for near-ties, we constructed the arborescence 100 times, each time using a Monte Carlo algorithm for selecting a high-consensus incoming edge for each node. Specifically, at every run of the arborescence algorithm, for each node *v* with a set of incoming edges E_i_, the consensus scores w_i_ were transformed as w’_i_ = w_i_ · u(0.95,1.05), where u is a continuous uniform distribution. An edge was selected for inclusion in the modified “jittered” arborescence if it achieved the maximum w’_i_ in at least one of the total 100 iterations. To effectively visualize the network, the *x*-coordinates of the layout were mapped to angular positions, transforming a hierarchical vertical depth into a radial projection.

Phosphosite level networks were created using the “subgraph” function from the R library *igraph*^103^, by including all phosphosites between proteins along edges from the jittered arborescence protein-level network. The “betweenness” centrality of network nodes was calculated by the “betweenness” *igraph* function using the inverse of edge consensus scores and corresponds roughly to the number of geodesics or shortest paths that must pass through a node.

### PHONEMeS network gene ontology annotation

To identify known protein functions, the full set of GO BP terms for all proteins present within the PHONEMeS network was obtained using *useEnsembl()* as implemented in the *biomaRt v2.62.1* R library^104,105^. To more broadly classify protein biological processes, GO BP terms were semantically clustered (**Supplementary Fig. 6**) using a binary cut algorithm with default parameters, as implemented in the *simplifyEnrichment v2.0.0* R library^106^, producing 22 mutually exclusive and distinct GO term clusters representing different biological processes. Of these, distinct clusters related to cytoskeletal organization, cell polarization, and adhesion were observed. GO terms within these clusters were further refined by manual curation to ensure that they were specifically related to migration-relevant processes. This dictionary of refined GO BP terms was used to annotate nodes of the PHONEMeS network.

## Supporting information

Supplementary File 1: Processed phosphoproteomic data and statistical analysis results

Supplementary File 2: Gene ontology overrepresentation analysis results for all significantly regulated proteins/genes

Supplementary File 3: Details for proteins annotated with cytoskeletal process-related GO terms

Supplementary File 4: Details for proteins annotated with nuclear process-related GO terms

Supplementary File 5: Manually added interaction edges

Supplementary File 6: Cytoscape network files

Supplementary File 7: Kinase inhibitor screening data

## Data availability

All raw phosphoproteomic data have been deposited at PRIDE with the ProteomeXchange dataset identifier [to be generated].

## Code availability

R scripts for mass spectrometry data analysis, R/Python scripts for network analysis, and a forked version of PHONEMeS are available at https://github.com/Kufalab-UCSD/CCR2-phospho/. IBM ILOG CPLEX Optimization Studio was used as an integer linear programming solver. The project-specific version of PHONEMeS was developed from the PHONEMeS-ILP source, under the GNU General Public License v3.0, available at https://github.com/saezlab/PHONEMeS-ILP.

## Acknowledgements

We thank: Weijun Xu and Enio Gjerga for help with the PHONEMeS software; the laboratories of Lan Nguyen and Roger Daly (Monash University) and Chris Langendorf (St Vincent’s Institute of Medical Research) for gifts of kinase inhibitors; and members of the Stone, Kufareva and Foster labs for productive discussions and critical review of the work. This work was supported by funding from National Health and Medical Research Council Project Grants APP1140867 and APP1140874 (M.J.S.), Ideas Grant APP2012579 (M.J.S., S.R.F., I.K.), and Investigator Grant 2025521 (R.P.B.); National Institute of Health grants R01 GM136202, R01 AI161880, and R01 AI118985 (T.M.H. and I.K.), and R21 AI149369 and R21 AI156662 (I.K.); University of California Office of the President (UCOP) Cancer Research Coordinating Committee (CRCC) seed grant C26CR10041 (I.K.); UCSD/UCSF Cancer Cell Mapping Initiative Pilot Grant under NIH U54 CA274502 (I.K.); and a Bioplatforms Australia ACvA Research Catalyst Program grant (S.R.F.).

## Author contributions

M.J.S., S.R.F., I.K., R.B.S. and T.M.H. conceived, designed and supervised the study. S.S.S., A.L., R.P., A.L.M., M.S., H.R., C.H., S.R.D., R.P.B., J.R.S., R.B.S., S.R.F., I.K. and M.J.S. performed experiments and analyzed results. A.L., S.S.S., T.Nguyen, W.C.C., T.Ngo, H.R., I.K. and M.J.S. wrote code for and performed the network modeling, analysis and presentation. S.S.S., A.L., M.S. and D.H.D. selected kinase inhibitors. S.S.S., A.L., R.P., S.R.F., I.K. and M.J.S. wrote the manuscript. All authors contributed to editing the manuscript.

## Competing interests

The authors declare no competing interests.

## Additional Information

**Supplementary information:** The online version contains supplementary material available at (URL to be determined).

**Correspondence** and requests for materials should be addressed to Martin Stone, Irina Kufareva or Simon Foster.

**Peer review information:** To be provided by journal.

**Reprints and permissions information** is available at (URL to be determined).

**Publisher’s note:** To be provided by journal.

**Open Access:** To be provided by journal.

## Supplementary Figures

**Supplementary Figure 1:** Chemokine receptor mRNA expression in THP-1 monocytes.

**Supplementary Figure 2:** CCL2 treatment induces time-dependent changes in protein phosphorylation

**Supplementary Figure 3:** Kinase Enrichment Analysis

**Supplementary Figure 4:** Clustering of phosphosite time courses

**Supplementary Figure 5:** Interplay between CCR2 and CCR1

**Supplementary Figure 6:** Semantic clustering of Gene Ontology Biological Processes by binary cut

**Supplementary Figure 7:** Divergent signaling pathways converge on the chemotaxis-related protein WASF2

**Supplementary Figure 8:** Primary kinase inhibitor screening

**Supplementary Figure 9:** Data quality control for phosphosite quant analysis on the CCL2 phosphoproteomics dataset

## Supplementary Tables

**Supplementary Table 1:** Regulators of cytoskeletal dynamics downstream of RAC1 and/or CDC42

**Supplementary Table 2:** Primers for qRT-PCR of chemokine receptors

## Supplementary Files

**Supplementary File 1:** Processed phosphoproteomic data and statistical analysis results

**Supplementary File 2:** Gene ontology overrepresentation analysis results for all significantly regulated proteins/genes

**Supplementary File 3:** Details for proteins annotated with cytoskeletal process-related GO terms

**Supplementary File 4:** Details for proteins annotated with nuclear process-related GO terms

**Supplementary File 5:** Manually added interaction edges

**Supplementary File 6:** Cytoscape networks, including: all network views presented in the manuscript; the full network with phosphosites; and the consensus-filtered network with phosphosites

**Supplementary File 7:** Kinase inhibitor screening data

## Footnotes

1 Square brackets indicate protein name abbreviations used in the PHONEMeS network, where they differ from commonly used abbreviations.

## Notes

### Competing Interest Statement

The authors have declared no competing interest.

