## Supplementary Figures and Tables for "Monocyte Migration Emerges from a Divergent Chemokine Signaling Network"

#### Supplementary Data

- Supplementary Figures 1-9
- Supplementary Tables 1-2

### Supplementary Figures

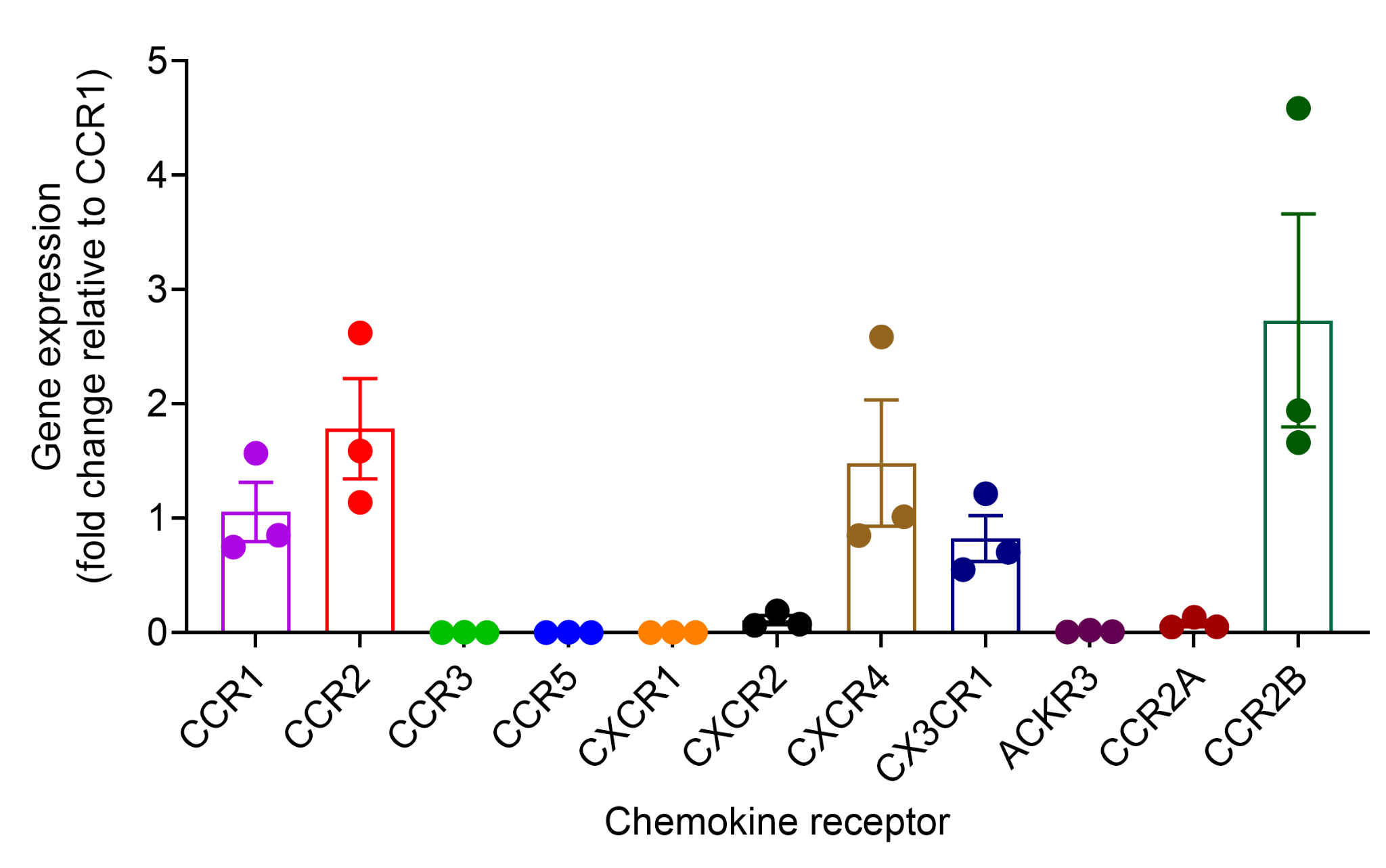

#### Supplementary Figure 1: Chemokine receptor mRNA expression in THP-1 monocytes.

Quantitative real-time PCR (qRT-PCR) was used to assess chemokine receptor gene expression of. Gene expression was calculated using the ΔΔC_t_ method and normalized to CCR1. Bars represent mean ± SEM of 3 independent experiments.

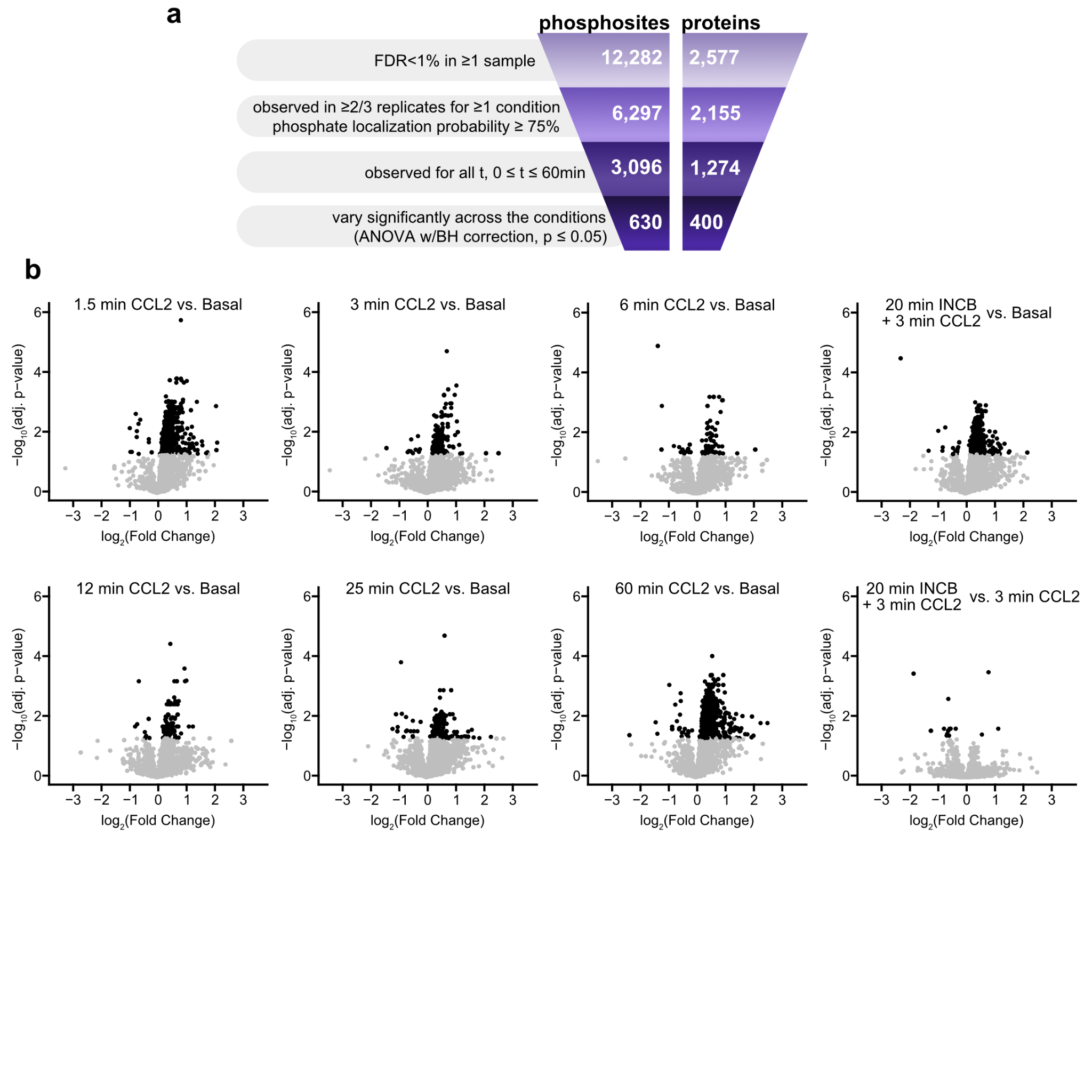

#### Supplementary Figure 2: CCL2 treatment induces time-dependent changes in protein phosphorylation.

**a,** A schematic indicating the number of phosphosites (left) and their proteins (right) detected at each phase of data processing, indicated on the left of the funnel. **b,** Volcano plots indicating relative changes in phosphosite intensity at various time points as indicated in the plot titles. Each point corresponds to a single phosphosite; those with a Benjamini-Hochberg post-hoc corrected p-value < 0.05 (adj. p-value) are colored black, whilst other points are colored gray.

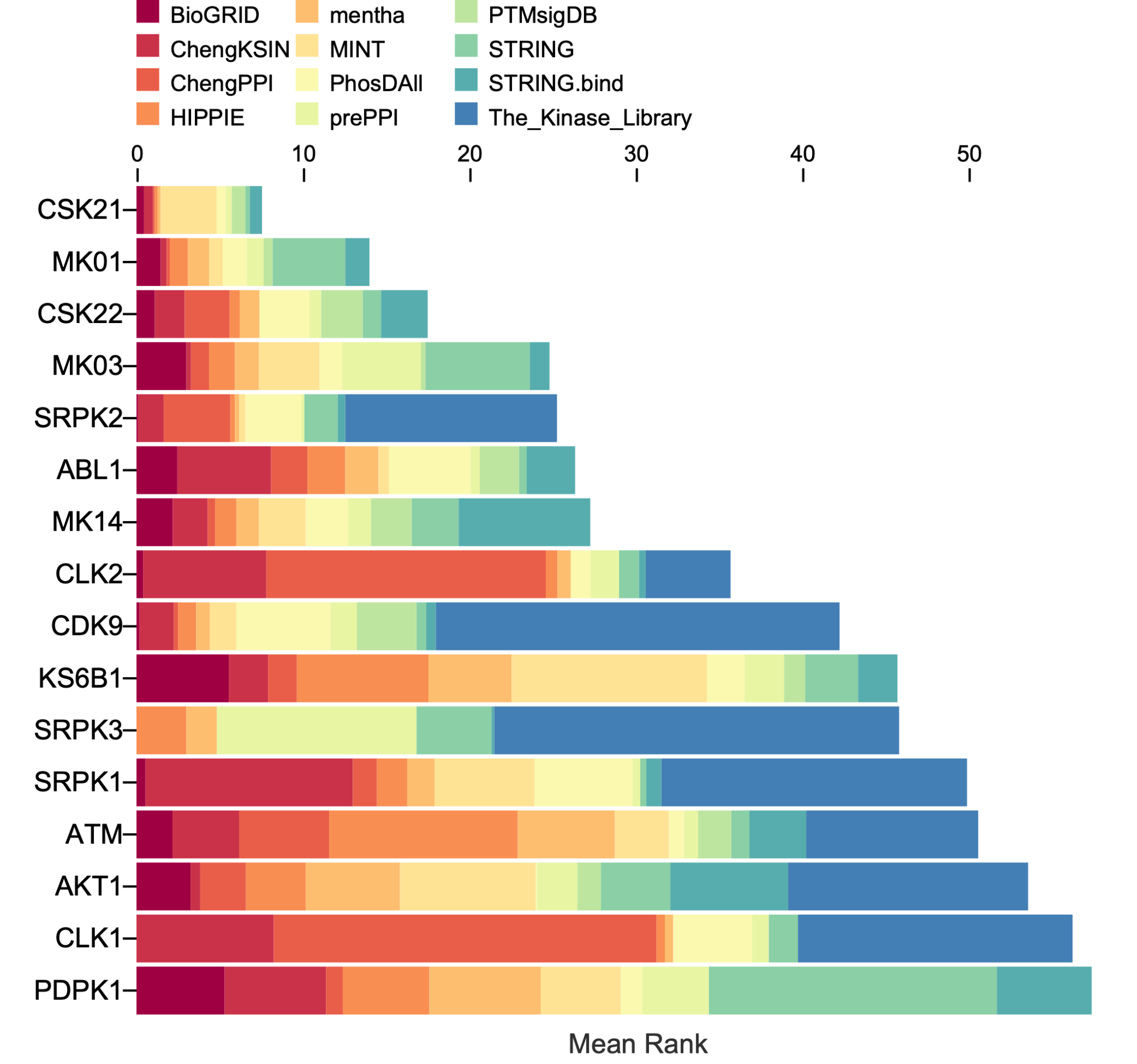

#### Supplementary Figure 3: Kinase Enrichment Analysis (KEA3) of the proteins that harbor the 630 phosphosites significantly regulated in CCL2-stimulated THP-1 cells.

Kinases are ranked based on the KEA3 “Mean Rank” score, which represents the average rank of the kinase across multiple libraries integrated into KEA3. A lower “Mean Rank” indicates a kinase that has been reported to phosphorylate a larger fraction of the 400 proteins submitted for the analysis (those harboring at least one of the significantly regulated phosphosites (ANOVA adj. p-value < 0.05)). Colored segments correspond to specific evidence libraries, including kinase-substrate interactions (e.g., PTMsigDB, PhosDAll), protein-protein interactions (e.g., BioGRID, STRING, MINT), and co-expression data. Results were generated using the KEA3 web interface (<https://maayanlab.cloud/kea3/>). (Kuleshov et al. Nucleic Acids Res., 2021: W304-W316. doi: 10.1093/nar/gkab359).

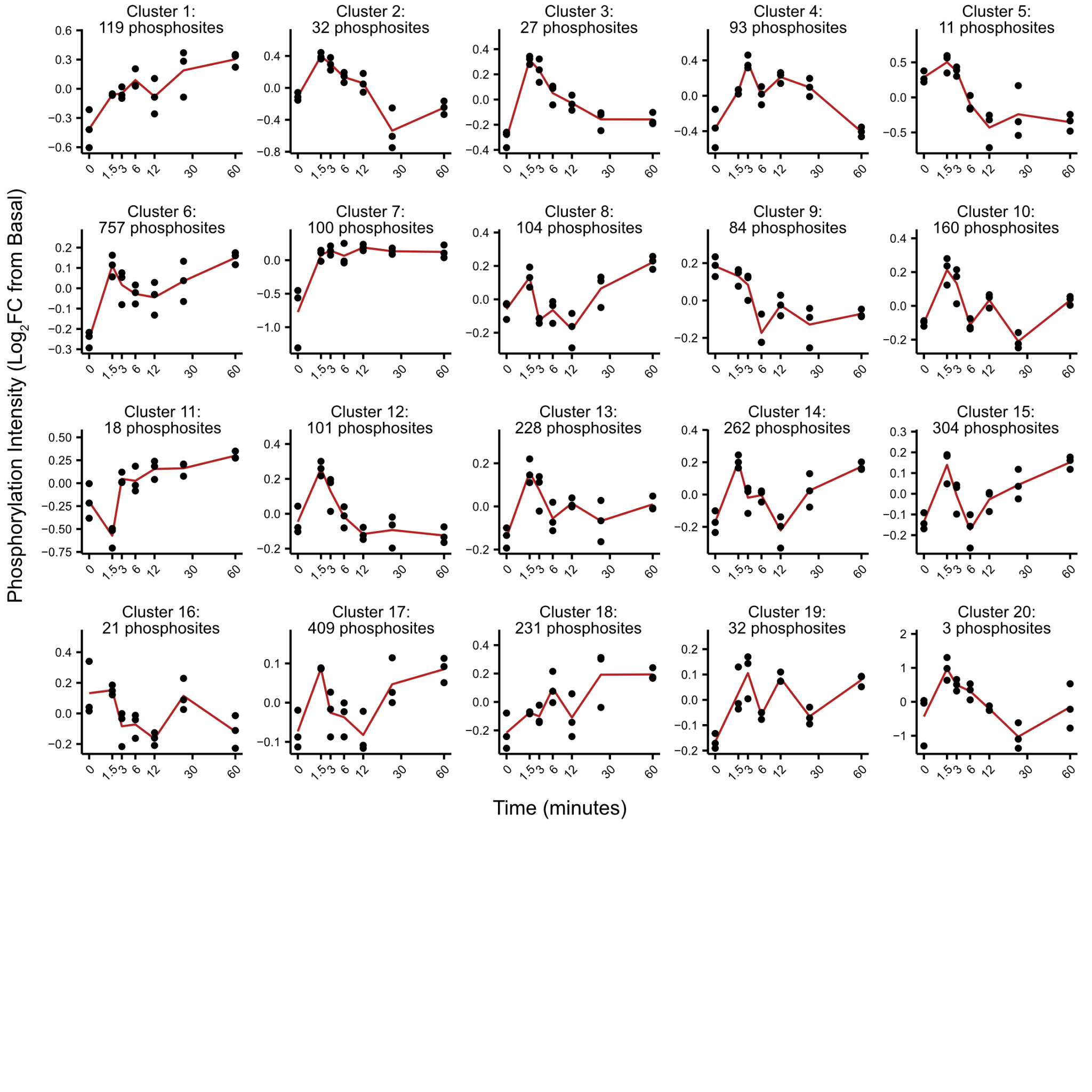

#### Supplementary Figure 4: Clustering of phosphosite time courses.

The time courses for 3,096 phosphosites detected at all time points in phosphoproteomics (**Fig. 1d**) were hierarchically clustered, as described in **Methods**. Clustering generated 20 time course clusters of varying size. Points represent averages of 3 replicates across all cluster members. The red line indicates the mean of the replicates. Time is represented on a square-root axis. For each plot, the header shows the cluster identifier and the number of members.

**a**

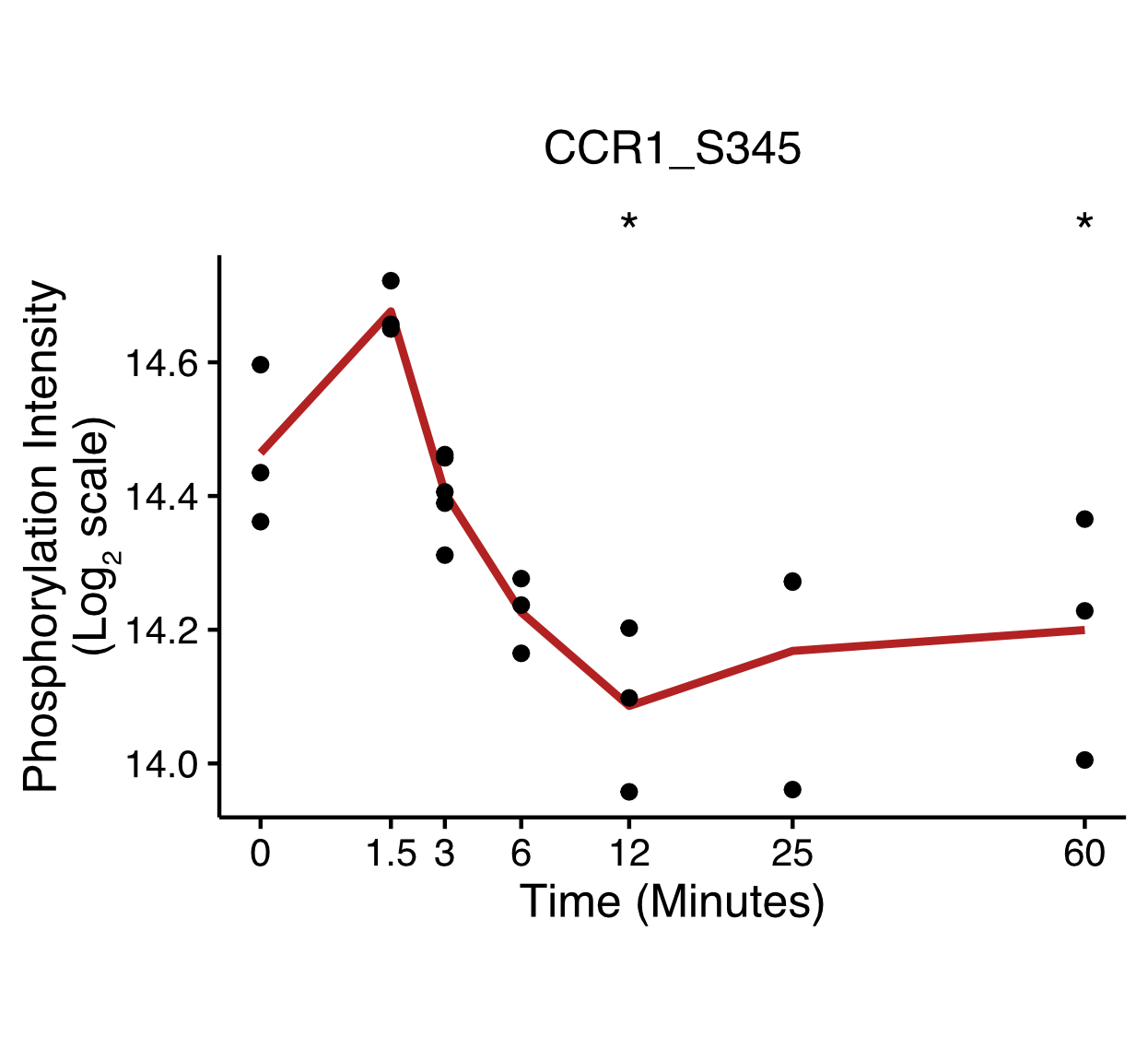

**b**

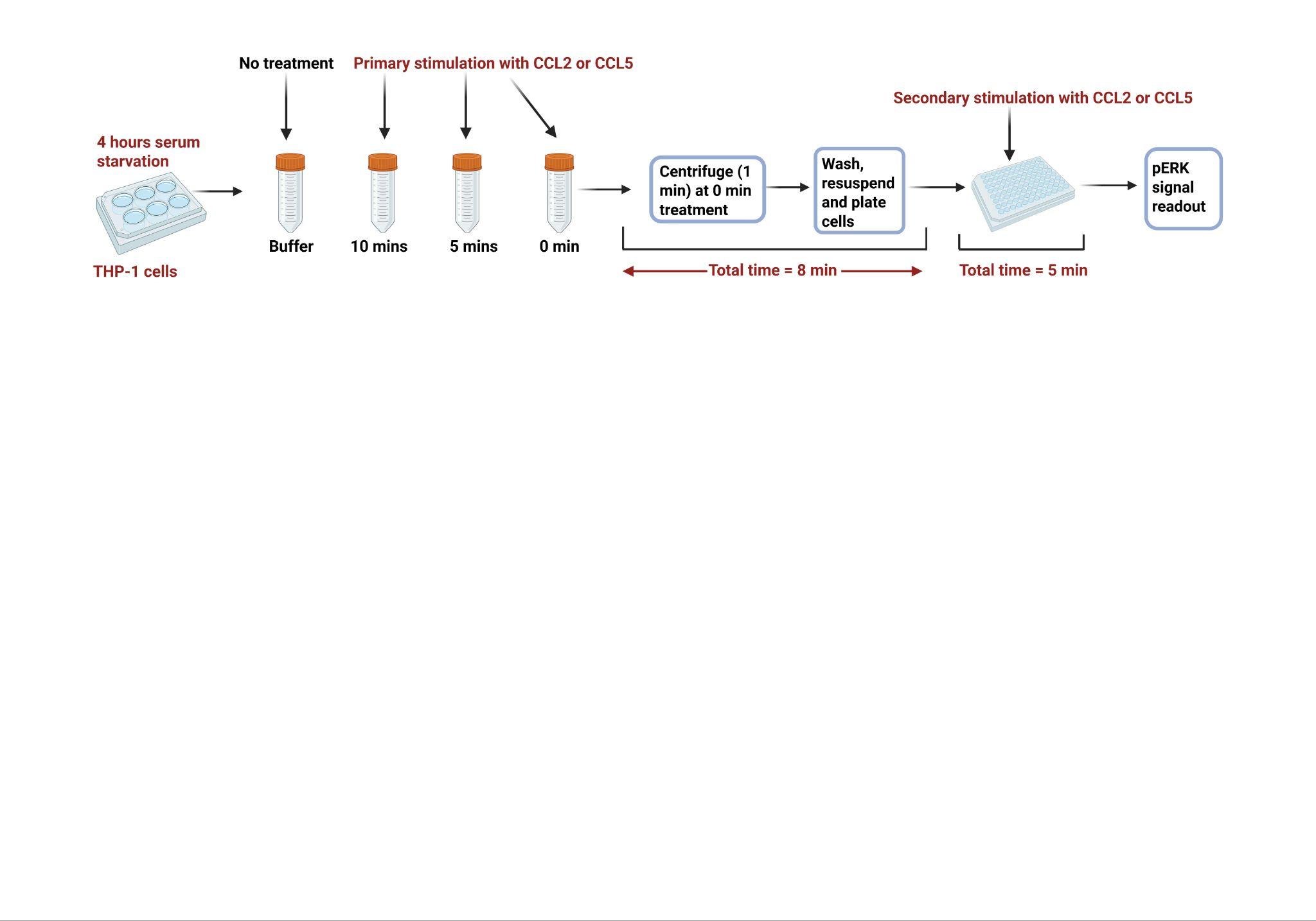

**c	d**

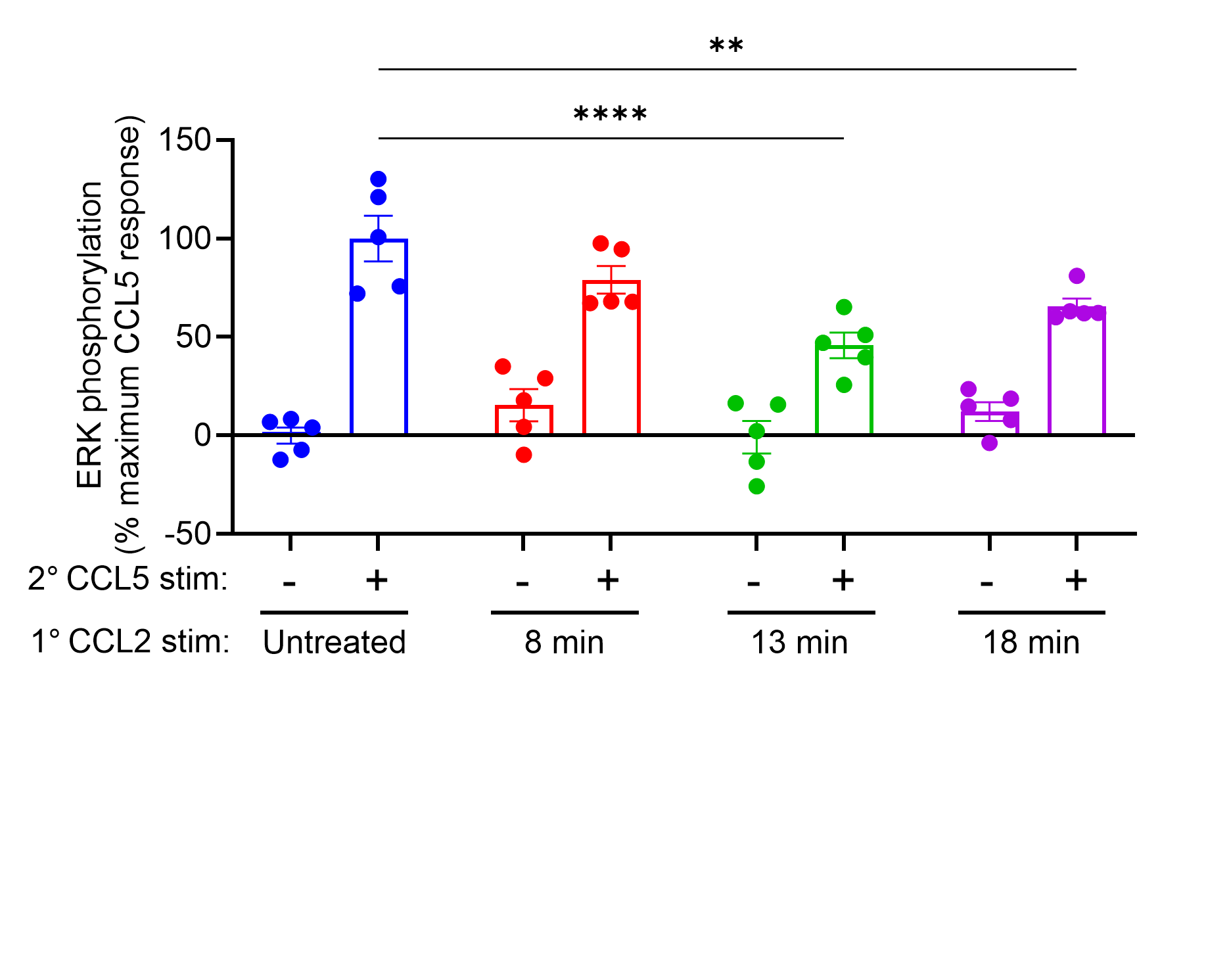

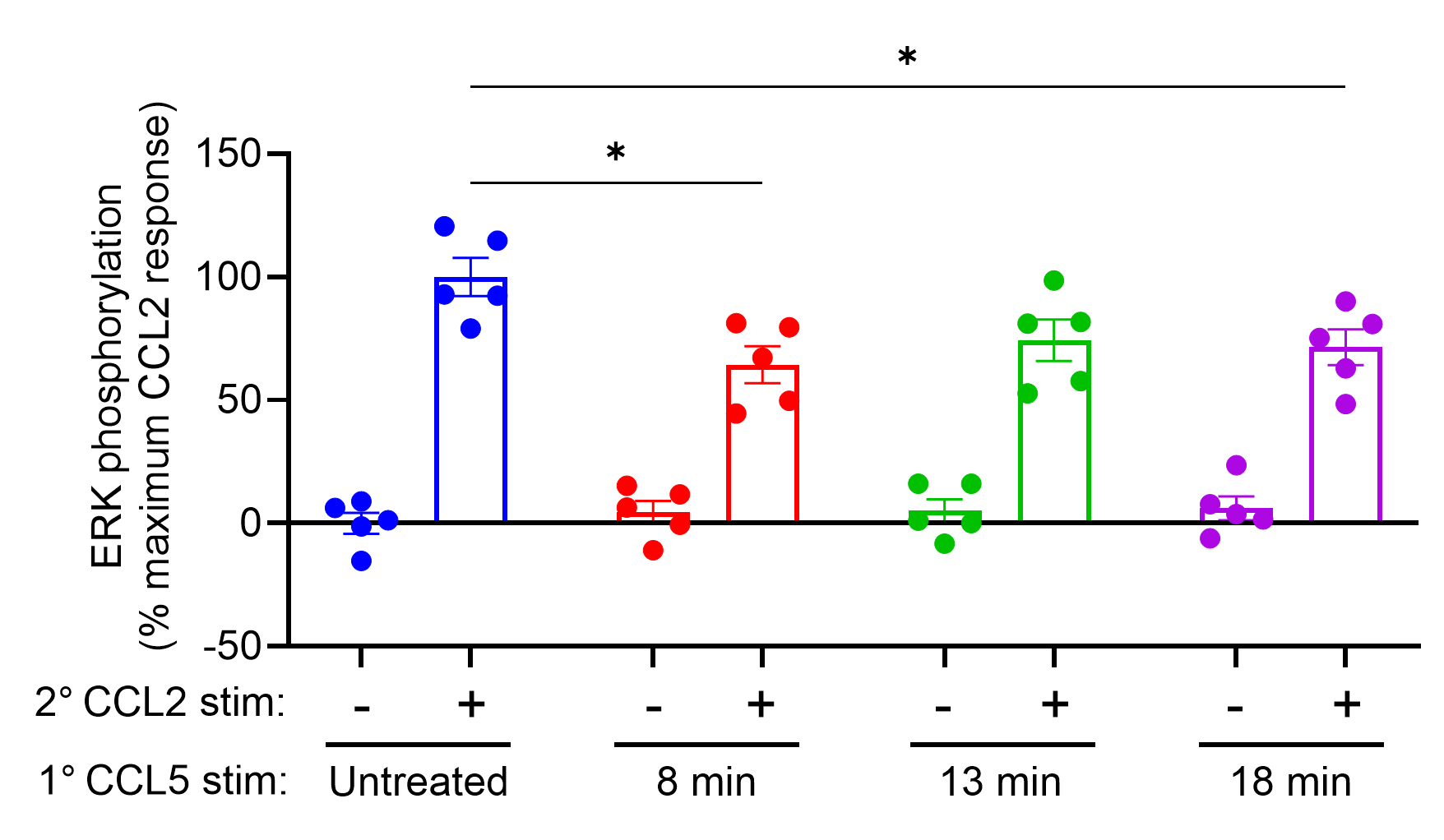

#### Supplementary Figure 5: Interplay between CCR2 and CCR1.

**a,** Global phosphoproteomics data show that CCL2 induced time-dependent changes in CCR1 phosphorylation in THP-1 monocytes, with phosphopeptide intensities (mean ± SEM, n=3) analyzed relative to baseline using one-way ANOVA with Benjamini-Hochberg correction (*p<0.05). **b,** Schematic diagram of CCR1/CCR2 heterologous desensitisation experiment. THP-1 monocytes were serum-starved for 4 h and then subjected to primary stimulation with CCL2 or CCL5 (100 nM) for 10, 5 or 0 min before centrifugation, while control cells received buffer. Cells were then washed, resuspended, plated and subjected to a secondary stimulation with CCL2 or CCL5 (100 nM) for 5 min; the total time elapsed between primary and secondary stimulation was 18, 13, or 8 min, respectively. Responses were measured by ERK phosphorylation (pERK). **c,** Heterologous desensitisation of CCR1 (CCL5 100 nM; secondary stimulation (2^0^)) following primary stimulation (1^0^) of CCR2 with CCL2 (100 nM). **d,**. Heterologous desensitisation of CCR2 (CCL2 100 nM; secondary stimulation (2^0^)) following primary stimulation (1^0^) of CCR1 with CCL5 (100 nM). ERK phosphorylation responses are expressed as a percentage of the maximal response to each chemokine (100 nM), derived from five independent experiments; statistical significance was determined using two-way ANOVA with Dunnett’s multiple comparisons test, comparing secondary CCL2 or CCL5 responses after primary stimulation to responses without primary stimulation (untreated control) (*p<0.05, **p<0.01, ***p<0.001).

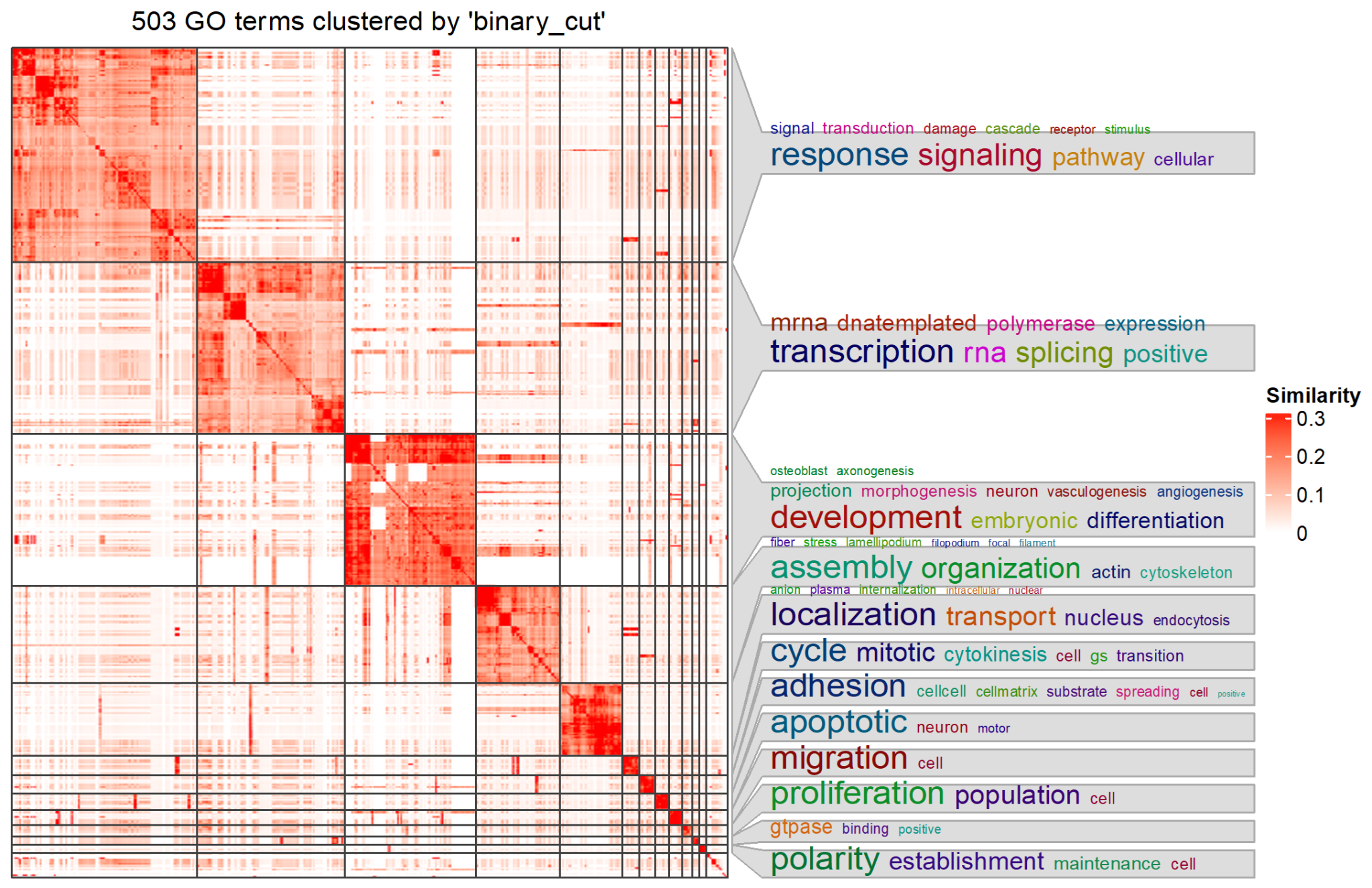

#### Supplementary Figure 6: Semantic clustering of Gene Ontology Biological Processes by binary cut.

The full set of all Gene Ontology Biological Process terms for all proteins presented in the PHONEMeS network were clustered by a binary cut algorithm as described in **Methods** and visualized as a similarity heatmap (white, least similar; red, most similar). The most prevalent terms within each cluster are shown in word clouds labeling each cluster.

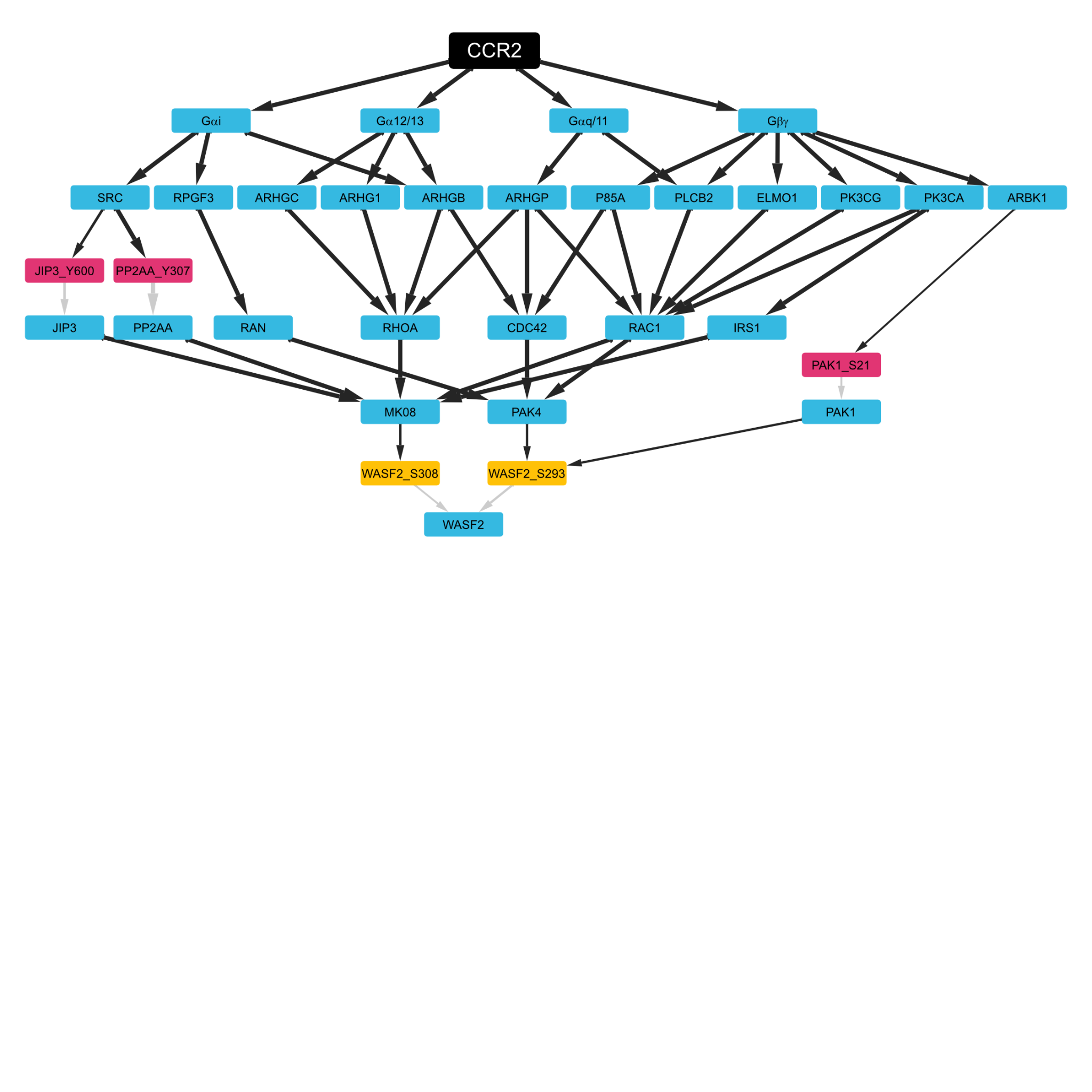

#### Supplementary Figure 7: Divergent signaling pathways converge on the chemotaxis-related protein WASF2.

An example of how divergent signals converge on chemotaxis-related proteins is shown for WASF2, a critical regulator of the transmission of signals from small GTPases to the actin cytoskeleton. For this subnetwork, the representation of the strongest consensus signaling cascade resulting in the phosphorylation of WASF2 is shown. Activation of WASF2 phosphosites is regulated by divergent branches of the CCR2 signaling network. Node colors: black, CCR2; blue, protein; rose, unobserved phosphosite; yellow, observed and significantly regulated phosphosite. Black arrows represent connections between proteins or from one protein to a phosphosite on another protein, whereas gray arrows indicate connections from a phosphosite to the protein on which it is located.

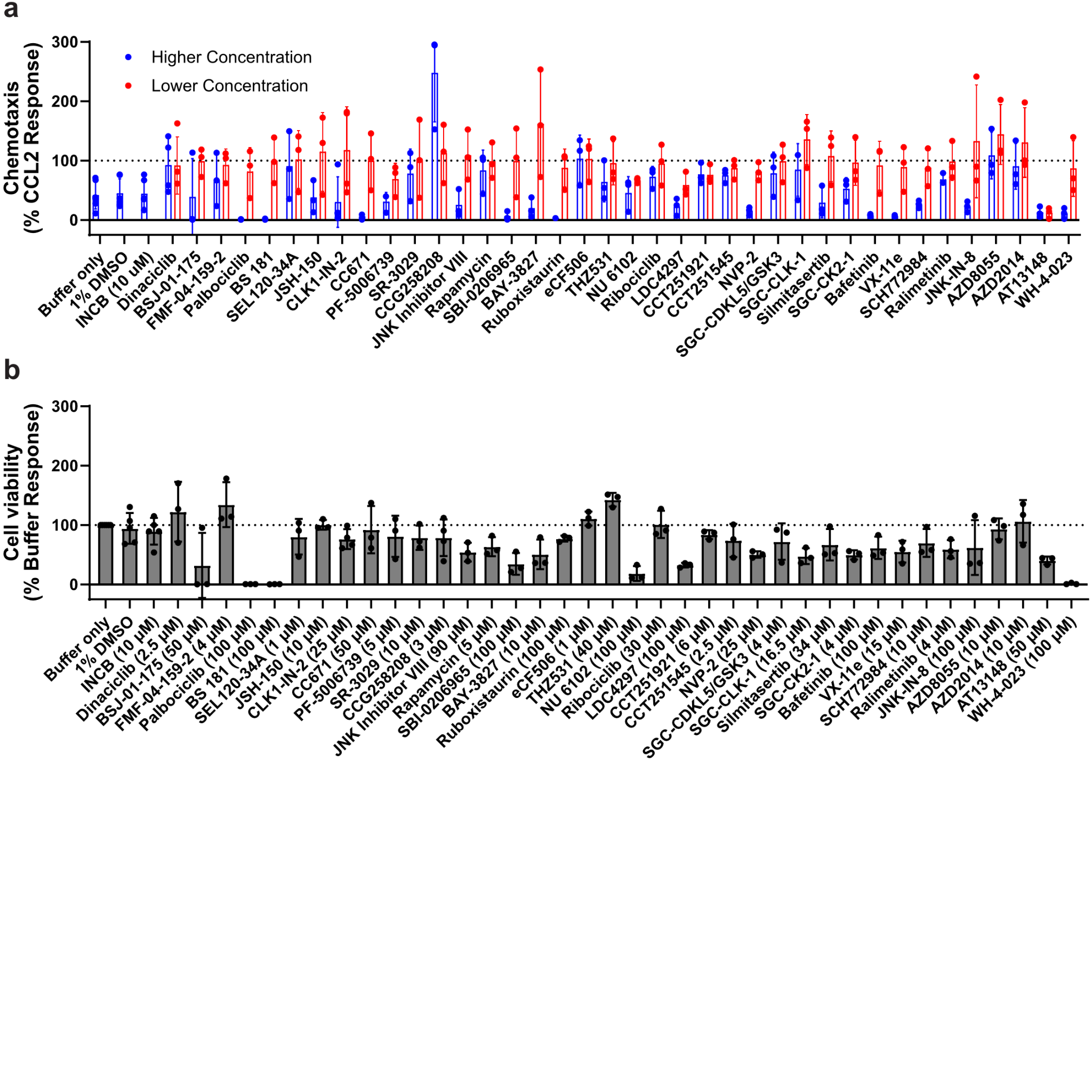

#### Supplementary Figure 8: Primary kinase inhibitor screening.

**a,b**, Kinase inhibitors were screened for their ability to inhibit THP-1 (**a**) chemotaxis and confounding effects on (**b**).cell viability THP-1 cells were pretreated for 2 hours with buffer, buffer with 1% DMSO, INCB3344, or two concentrations of kinase inhibitor with a 100-fold difference (higher, blue; lower, red). For chemotaxis, pretreated THP-1 cells were stimulated with 10 nM CCL2. Cell viability was assessed in parallel over the duration of the chemotaxis assay (4 hours total) using the higher concentration for each inhibitor or pretreatment controls. Detailed concentrations used for each inhibitor are available in **Supplementary File 7**. Points and error bars represent the means and standard deviations for n=3-4 experiments for inhibitors (n=6 for controls), performed in triplicate.

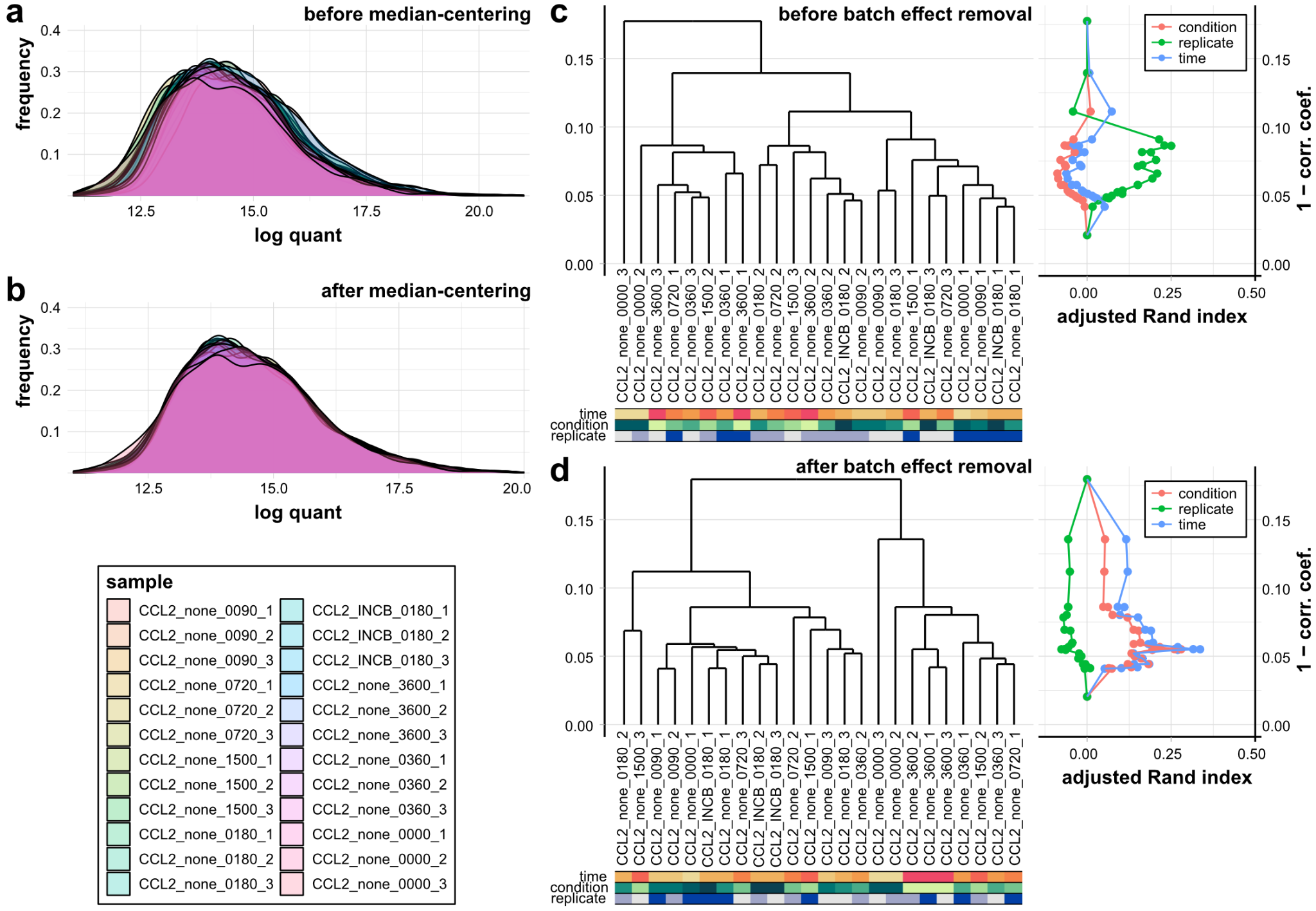

#### Supplementary Figure 9: Data quality control for phosphosite quant analysis on the CCL2 phosphoproteomics dataset.

**a,b**, Peptide-spectrum match quant distribution for all samples before (**a**) and after (**b**) median centering. **c,d**, Adjusted Rand indices between the sample-to-sample correlation dendrogram cut at different heights and the experimental variables were compared before (**c**) and after (**d**) batch effect removal. Sample names are comprised of four terms separated by underscores. The first indicates the chemokine treatment; the second indicates the pretreatment; the third indicates the time point in seconds; the fourth indicates the replicate of the sample.

### Supplementary Tables

#### Supplementary Table 1: Regulators of cytoskeletal dynamics downstream of RAC1 and/or CDC42

| Protein | Function |
| --- | --- |
| WASF2 | Binds ARP2/3 to stimulate nucleation of actin filaments |
| NCK1 | Activator of the WASP/ARP2/3 complex |
| ITPI2 | Actin filament binding protein |
| MYO18 [MY18A] | Promotes retrograde actin-myosin flow required for protrusion |
| ACAP2 | Contributes to cytoskeletal rearrangement and stress fiber formation |
| CLAP1 | Stabilizes microtubules at the leading edge |
| ADDA | Promotes assembly of the spectrin-actin network linking the cytoskeleton to the plasma membrane |
| PKHA2 | Promotes cell adhesion to the extracellular matrix |

#### Supplementary Table 2: Primers for qRT-PCR of chemokine receptors

| Target Gene | NCBI Refseq | For/Rev | Primer Sequence (5' > 3') | Primer Length | Tm | Location in ORF | Amplicon Length |
| --- | --- | --- | --- | --- | --- | --- | --- |
| CCR1 | NM_001295 | FOR Seq | GACTATGACACGACCACAGAGT | 22 | 61.1 | 25-46 | 128 |
|  |  | REV seq | CCAACCAGGCCAATGACAAATA | 22 | 60.8 | 152-131 | 128 |
| CCR2 | NM_001123396 | FOR Seq | TACGGTGCTCCCTGTCATAAA | 21 | 60.6 | 82-102 | 120 |
|  |  | REV seq | TAAGATGAGGACGACCAGCAT | 21 | 60.4 | 201-181 | 120 |
| CCR3 | NM_178329 | FOR Seq | TGGCATGTGTAAGCTCCTCTC | 21 | 61.5 | 309-329 | 85 |
|  |  | REV seq | CCTGTCGATTGTCAGCAGGATTA | 23 | 62 | 393-371 | 85 |
| CCR5 | NM_000579 | FOR Seq | TTCTGGGCTCCCTACAACATT | 21 | 60.8 | 739-759 | 93 |
|  |  | REV seq | TTGGTCCAACCTGTTAGAGCTA | 22 | 60.4 | 831-810 | 93 |
| CXCR1 | NM_000634 | FOR Seq | CTGACCCAGAAGCGTCACTTG | 21 | 62.9 | 436-456 | 139 |
|  |  | REV seq | CCAGGACCTCATAGCAAACTG | 21 | 60.1 | 574-554 | 139 |
| CXCR2 | NM_001557 | FOR Seq | CCTGTCTTACTTTTCCGAAGGAC | 23 | 60.5 | 535-557 | 82 |
|  |  | REV seq | TTGCTGTATTGTTGCCCATGT | 21 | 60.5 | 616-596 | 82 |
| CXCR4 | NM_003467 | FOR Seq | ACTACACCGAGGAAATGGGCT | 21 | 63 | 32-52 | 133 |
|  |  | REV seq | CCCACAATGCCAGTTAAGAAGA | 22 | 60.2 | 164-143 | 133 |
| CX3CR1 | NM_001171174 | FOR Seq | AGTGTCACCGACATTTACCTCC | 22 | 61.7 | 286-307 | 134 |
|  |  | REV seq | AAGGCGGTAGTGAATTTGCAC | 21 | 61.2 | 419-399 | 134 |
| ACKR3 | NM_020311 | FOR Seq | TCTGCATCTCTTCGACTACTCA | 22 | 60 | 6-27 | 130 |
|  |  | REV seq | GTAGAGCAGGACGCTTTTGTT | 21 | 60.6 | 135-115 | 130 |
| CCR2a | NM_001123041.2 | FOR Seq | GCTGCATCAATCCCATCATCTA | 22 | 62 | 1378-1400 | 81 |
|  |  | REV seq | CAATCCTACAGCCAAGAGCTATG | 23 | 63 | 1436-1459 | 81 |
| CCR2b | NM_001123396.2 | FOR Seq | TGAATGGGAGTGAGGGATAGT | 21 | 62 | 1939-1960 (3' UTR) | 96 |
|  |  | REV seq | CCTTTGCTCACCTTTGTCTTTG | 22 | 62 | 2013-2035 | 96 |
| 18S | NR_146146.1 | FOR seq | TCGAGGCCCTGTAATTGGAA | 20 | 57 | n/a | 61 |
|  |  | REV seq | CCCTCCAATGGATCCTCGTT | 20 | 58 | n/a | 61 |
